# A Framework for Designing Efficient Deep Learning-Based Genomic Basecallers

**DOI:** 10.1101/2022.11.20.517297

**Authors:** Gagandeep Singh, Mohammed Alser, Kristof Denolf, Can Firtina, Alireza Khodamoradi, Meryem Banu Cavlak, Henk Corporaal, Onur Mutlu

## Abstract

Nanopore sequencing generates noisy electrical signals that need to be converted into a standard string of DNA nucleotide bases using a computational step called basecalling. The performance of basecalling has critical implications for all later steps in genome analysis. Therefore, there is a need to reduce the computation and memory cost of basecalling while maintaining accuracy. We present `RUBICON`, a framework to develop efficient hardware-optimized basecallers. We demonstrate the effectiveness of `RUBICON` by developing `RUBICALL`, the first hardware-optimized mixed-precision basecaller that performs efficient basecalling, outperforming the state-of-the-art basecallers. We believe `RUBICON` offers a promising path to develop future hardware-optimized basecallers.

## 1. Background

The rapid advancement of genomics and sequencing technologies continuously calls for the adjustment of existing algorithmic techniques or the development of entirely new computational methods across diverse biomedical domains [1–14]. Modern sequencing machines [15, 16] are capable of sequencing complex genomic structures and variants with high accuracy and throughput using long-read sequencing technology [17]. Oxford Nanopore Technologies (ONT) is the most widely used long-read sequencing technology [17–22]. ONT devices generate long genomic reads, each of which has a length ranging from a few hundred to a million base pairs or nucleotides, i.e., A, C, G, and T in the DNA alphabet [23–27].

ONT devices sequence a genome by measuring changes to an electrical signal as a single strand of DNA is passed through a nanoscale hole or *nanopore* [28]. The generated noisy electrical signal or *squiggle* is decoded into a sequence of nucleotides using a computationally-expensive step, called *basecalling* [19, 29–32]. Basecallers need to address two key challenges to accurately basecall a raw sequencing input. First, providing accurate predictions of each and every individual nucleotide, as the sensors measuring the changes in electrical current can only measure the effect of multiple neighboring nucleotides together [29]. Second, tolerating low signal-to-noise ratio (SNR) caused by thermal noise and the lack of statistically significant current signals triggered by DNA strand motions [30].

Modern basecallers use deep learning-based models to significantly (by at least 10%) improve the accuracy of predicting a nucleotide base from the squiggle compared to traditional non-deep learning-based basecallers [16–18, 31, 33–37]. The success of deep learning in genome basecalling is attributed to the advances in its architecture to model and identify spatial features in raw input data to predict nucleotides.

However, we observe the following six shortcomings with the current basecallers [33, 38–45]. First, current state-of-the-art basecallers are slow and show poor performance on state-of-the-art CPU and GPU-based systems, bottlenecking the entire genomic analyses. For example, state-of-the-art throughput optimized basecaller, `Dorado-fast`, takes ∼ 2.1 hours to basecall a 300 Gbps (Giga basepairs) human genome at 3× coverage on a server-grade GPU (NVIDIA A10G [46] GPU with 24GiB DRAM and 16 ×CPU with 64 GiB DRAM) [47], while the subsequent step, i.e., read mapping, takes only a small fraction of basecalling time (∼ 0.11 hours using minimap2 [48]). We observe that basecalling is the single longest stage in the genome sequencing pipeline, taking up to 43% of execution time while the subsequent step of overlap finding, assembly, read mapping, and polishing take 18%, 4%, <1%, and 35% of execution time, respectively.

Second, for real-time sequencing, high basecalling throughput is a critical factor [7]. In particular, scenarios such as *field sequencing* [40] and *adaptive sampling* [49] necessitate rapid basecalling due to hardware limitations and the need for real-time decision-making. Field sequencing, often conducted in remote or resource-constrained environments, demands immediate basecalling to obtain actionable genomic information swiftly. Conventional high-compute infrastructure is often unavailable or impractical in these settings, underscoring the importance of an efficient basecalling process. Similarly, adaptive sampling protocols, aiming to optimize sequencing output based on real-time analysis of initial sequencing data, require a fast and accurate basecaller to make prompt decisions regarding read continuation or rejection. Also, enhancing the speed and efficiency of basecalling is critical for re-basecalling existing datasets using advanced, higher-accuracy models. By revisiting earlier data with improved basecalling algorithms, researchers can achieve a more precise representation of the genomic sequence. Current basecallers provide a tradeoff between speed and accuracy, often leading to sub-optimal performance in real-time sequencing scenarios.

Third, since basecalling shares similarities with automatic-speech recognition (ASR) task, many researchers have directly adapted established ASR models, such as Quartznet [50], Citrinet [51], and Conformers [52], for basecalling without customizing the neural network architecture specifically for the basecalling problem. Such an approach might lead to higher basecalling accuracy but at the cost of large and unoptimized neural network architecture. For example, `Bonito_CTC`, an expert-designed convolutional neural network (CNN)-based version of `Bonito` from ONT, has ∼ 10 million model parameters. We show in Section 2.1.1 that we can eliminate up to 85% of the model parameters to achieve a 6.67× reduction in model size without any loss in basecalling accuracy. Therefore, current basecalling models are costly to run, and the inference latency becomes a major bottleneck.

Fourth, modern basecallers are typically composed of convolution layers with skip connections^1^ [53] (allow reusing of activations from previous layers) that creates two major performance issues: (a) skip connections increase the data lifetime: the layers whose activations are reused in future layers must either wait for this reuse to occur before accepting new input or store the activations for later use by utilizing more memory. Thus, leading to high resource and storage requirements; and (b) skip connections often need to perform additional computation to match the channel size at the input of the non-consecutive layer, which increases the number of model parameters; e.g., `Bonito_CTC` requires ∼ 21.7% additional model parameters due to the skip connections.

Fifth, current basecallers use floating-point precision (32 bits) to represent each neural network layer present in a basecaller. This leads to high bandwidth and processing demands [54–56]. Thus, current basecallers with floating-point arithmetic precision have inefficient hardware implementations. We observe in Section 2.1.2 that the arithmetic precision requirements of current basecallers can be reduced ∼ 4× by adjusting the precision for each neural network layer based on the target hardware and desired accuracy. Sixth, basecallers that provide higher throughput have lower basecalling accuracy. For example, we show in Section 2.2 and Supplementary S4 that `Bonito_CRF-fast` provides up to 51.65× higher basecalling performance using 36.96× fewer model parameters at the expense of the 5.37% lower basecalling accuracy compared to most accurate basecaller.

These six problems concurrently make basecalling slow, inefficient, and memory-hungry, bottlenecking all genomic analyses that depend on it. Therefore, there is a need to reduce the computation and memory cost of basecalling while maintaining their performance. However, developing a basecaller that can provide fast runtime performance with high accuracy requires a deep understanding of genome sequencing, machine learning, and hardware design. At present, computational biologists spend significant time and effort to design and implement new basecallers by an extensive trial-and-error process.

**Our goal** is to overcome the above issues by developing a comprehensive framework for specializing and optimizing a deep learning-based basecaller that provides high efficiency and performance.

To this end, we introduce `RUBICON`, the first framework for specializing and optimizing a machine learning-based basecaller. `RUBICON` uses two machine learning techniques to develop hardware-optimized basecallers that are specifically designed for basecalling. First, we propose `QABAS`, a quantization-aware basecalling architecture search framework to specialize basecaller architectures for hardware implementation while considering hardware performance metrics (e.g., latency, throughput, etc.). `QABAS` uses neural architecture search (NAS) [57] to evaluate millions of different basecaller architectures. As discussed in Supplementary Section S1, during the basecaller neural architecture search, `QABAS` quantizes the neural network model by exploring and finding the best bit-width precision for each neural network layer, which largely reduces the memory and computational complexity of a basecaller. Adding quantization to the basecaller neural architecture search dramatically increases the model search space (∼ 6.72 ×10^20^ more viable options in our search space). However, jointly optimizing basecalling neural network architecture search and quantization allows us to develop accurate basecaller architectures that are optimized for hardware acceleration. Second, we develop `SkipClip` to remove all the skip connections present in modern basecallers to reduce resource and storage requirements without any loss in basecalling accuracy. `SkipClip` performs a skip removal process using knowledge distillation [58], as shown in Supplementary Figure S2 in Supplementary Section S2, where we train a smaller network (*student*) without skip connections to mimic a pre-trained larger network (*teacher*) with skip connections. Figure 1 shows the key components of `RUBICON`. It consists of four modules. `QABAS` 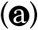 and `SkipClip` 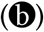 are two novel techniques that are specifically designed for specializing and optimizing machine learning-based basecallers. `RUBICON` provides support for `Pruning` 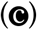, which is a popular model compression technique where we discard network connections that are unimportant to neural network performance [59–62]. We integrate `Training` 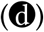 module from the official ONT basecalling pipeline [63]. For both the `Pruning` and `Training` modules, we provide the capability to use knowledge distillation [58, 64] for faster convergence and to increase the accuracy of the designed basecalling network.

**Figure 1:**
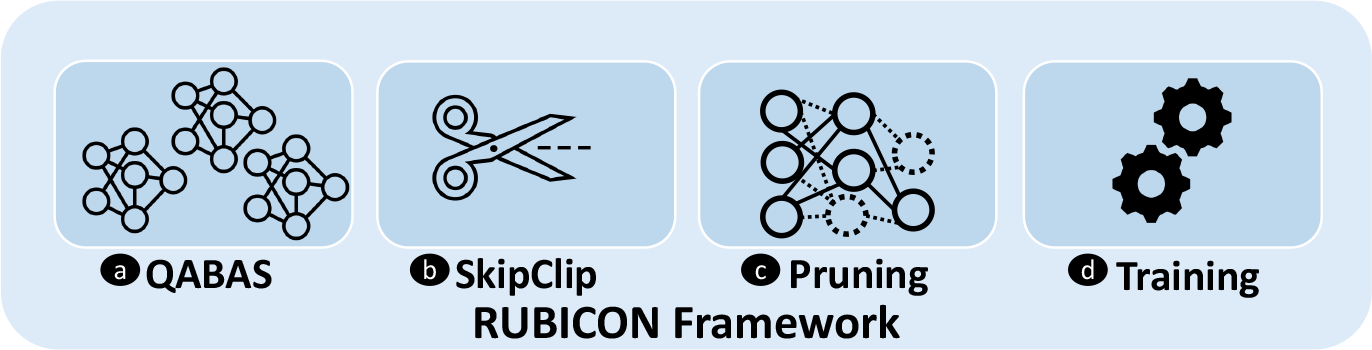
Overview of RUBICON framework.

### Key results

We demonstrate the effectiveness of `RUBICON` by developing `RUBICALL`, the first hardware-optimized mixed-precision basecaller that performs efficient basecalling, outperforming the state-of-the-art basecallers. Supplementary Figure S5 in Supplementary Section S2 shows the `RUBICALL` architecture. We compare `RUBICALL` to five different basecallers. We demonstrate six key results. First, `RUBICALL` provides, on average, 2.85% higher basecalling accuracy with 3.77× higher basecalling throughput compared to the fastest basecaller. Compared to an expert-designed basecaller `RUBICALL` provides 128.13× higher basecalling throughput without any loss in basecalling accuracy by leveraging mixed precision computation when implemented on a cutting-edge spatial vector computing system, i.e., the AMD-Xilinx Versal AIE-ML [65]. Second, we show that `QABAS`-designed models are 5.74× smaller in size with 2.41× fewer neural network model parameters than an expert-designed basecaller. Third, by further using our `SkipClip` approach, `RUBICALL` achieves a 6.88× and 2.94× reduction in neural network model size and the number of parameters, respectively. Fourth, we show in Supplementary S4 that compared to the most accurate state-of-the-art basecaller (i.e., `Bonito_CRF-sup`), `RUBICALL` provides 185.54 speedup using 19.22× lower parameters at the expense of, on average, 2.47% lower accuracy. Fifth, assemblies constructed using reads basecalled by `RUBICALL` lead to higher quality, more contiguous, and more complete assemblies for all evaluated species than that provided by other basecallers. Sixth, `RUBICALL` provides a 1.82%-26.49% lower number of base mismatches with the largest number of mapped bases and mapped reads compared to the baseline basecaller. Our experimental results on state-of-the-art computing systems show that `RUBICALL` is a fast, memory-efficient, and hardware-friendly basecaller. `RUBICON` can help researchers develop hardware-optimized basecallers that are superior to expert-designed models and can inspire independent future ideas.

## 2. Results

### 2.1. Analyzing the State-of-the-Art Basecaller

We observe established automatic-speech recognition (ASR) models being directly applied to basecalling without optimizing it for basecalling. Such an approach leads to large and unoptimized basecaller architectures. We evaluate the effect of using two popular model compression techniques on the `Bonito_CTC` basecaller: (1) Pruning, and (2) Quantization.

#### 2.1.1. Effect of Pruning

We show the effect of pruning `Bonito_CTC` on the validation accuracy and model size in Figure 2(a) and Figure 2(b), respectively. Pruning is a model compression technique where we discard network connections that are unimportant to network performance without affecting the inference accuracy [59–62]. We use unstructured element pruning and structured channel pruning with different degrees of sparsity. Unstructured or element pruning is a fine-grain way of pruning individual weights in a neural network without applying any pruning constraints. While in structured pruning, we remove a larger set of weights while maintaining a dense structure of the model [66, 67].

**Figure 2:**
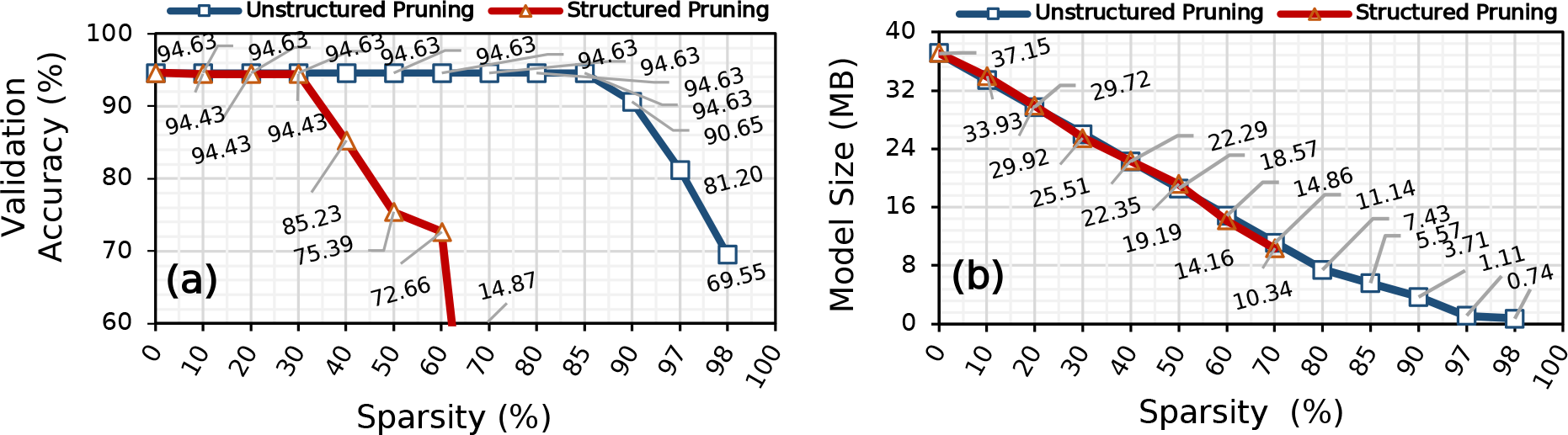
Effect of pruning the elements and channels of Bonito_CTC using unstructured and structured pruning, respectively, on: (a) validation accuracy and (b) model size.

We make three major observations. First, pruning up to 85% of the `Bonito_CTC` model weights using unstructured pruning reduces the model size by 6.67× while maintaining the same accuracy as the baseline, unpruned `Bonito_CTC` model. Unstructured pruning leads to the highest model compression [68] at the cost of having sparse weights structure that is unsuitable for acceleration on any hardware platform. While pruning 30-40% of the `Bonito_CTC` model filters, using structured pruning reduces the model size by 1.46-1.66× while maintaining the same accuracy of the baseline, unpruned `Bonito_CTC` model. Such a high pruning ratio shows that most of the weights are redundant and do not contribute to the actual accuracy. Second, after pruning 97% (60%) of the model weights, `Bonito_CTC` provides 81.20% (72.66%) basecalling accuracy while using 33.33× (2.62×) smaller model using unstructured pruning (structured pruning). Third, the *knee point*^2^ for unstructured pruning and structured pruning is at 98% and 60% where `Bonito_CTC` provides 65.14% and 72.66% of basecalling accuracy, respectively. Beyond the knee-point, `Bonito_CTC` losses its complete prediction power. We conclude that `Bonito_CTC` is over-parameterized and contains redundant logic and features.

#### 2.1.2. Effect of Quantization

Figure 3 shows the effect of using a quantized model to basecall on the basecalling accuracy for four different species. In Figure 4, we show the effect of quantization on the model size. We quantize both the weight and activation using six different bit-width configurations (*<*`3,2`*>, <*`4,2`*>, <*`4,4`*>, <*`4,8`*>, <*`8,4`*>*, and *<*`16,16`*>*). We also show the results with the default floating-point precision (*<*`fp32,fp32`*>*). We use static quantization that uses the same precision for each neural network layer.

**Figure 3:**
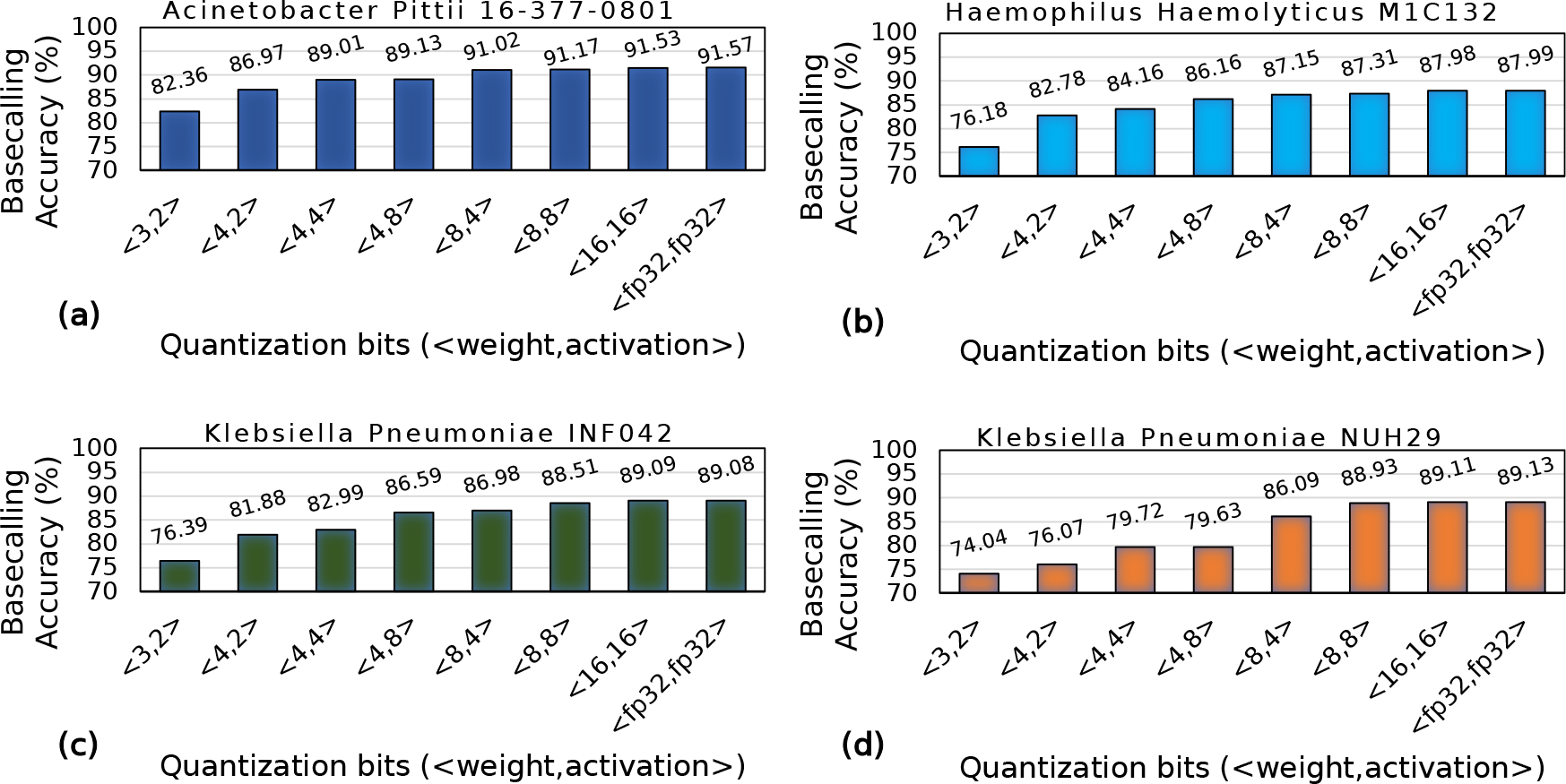
Basecalling using quantized models.

**Figure 4:**
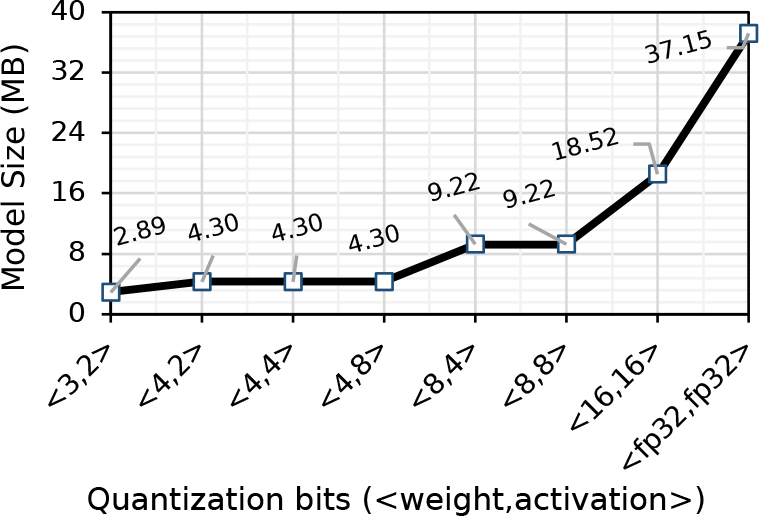
Effect of quantizing weight and activation of Bonito_CTC on model size. We quantize both the weight and activation with static precision. Since weights are the trainable parameters in a neural network, only weights contribute to the final model size.

We make four main observations. First, using a precision of *<*`8,8`*>* for weight and activation for all the layers of `Bonito_CTC` causes a negligible accuracy loss (0.18%-0.67%), while reducing the model size by 4.03×. Second, `Bonito_CTC` is more sensitive to weight precision than activation precision. For example, we observe a loss of 1.82%-9.48% accuracy when using a precision of *<*`4,8`*>* instead of *<*`16,16`*>* bits compared to an accuracy loss of only 0.51%-3.02% when using a precision of *<*`8,4`*>* instead of *<*`16,16`*>* bits. Third, we observe a significant drop in accuracy (by 9.17%-15.07%), when using less than 4 bits for weights (e.g., using *<*`3,2`*>* configuration). Fourth, using bit-width precision of *<*16,16*>* bits provides ∼2× reductions in model size and without any accuracy loss compared to using full precision (*<*`fp32,fp32`*>*) floating-point implementation. We conclude that the current state-of-the-art basecaller, `Bonito_CTC`, can still efficiently perform basecalling even when using lower precision for both the weight and activation.

### 2.2. RUBICALL: Overall Trend

We compare the overall basecalling throughput of `RUBICALL` with that of the baseline basecallers in terms of average basecalling accuracy, model parameters, and model size in Figure 5(a), 5(b), and 5(c), respectively. We evaluate `RUBICALL` using: (1) MI210 GPU [69] (`RUBICALL`-FP) using floating-point precision computation, and (2) Versal ACAP VC2802 [65], a cutting-edge spatial vector computing system (`RUBICALL`-MP) using mixed-precision computation. Section 5 provides details on our evaluation methodology.

**Figure 5:**
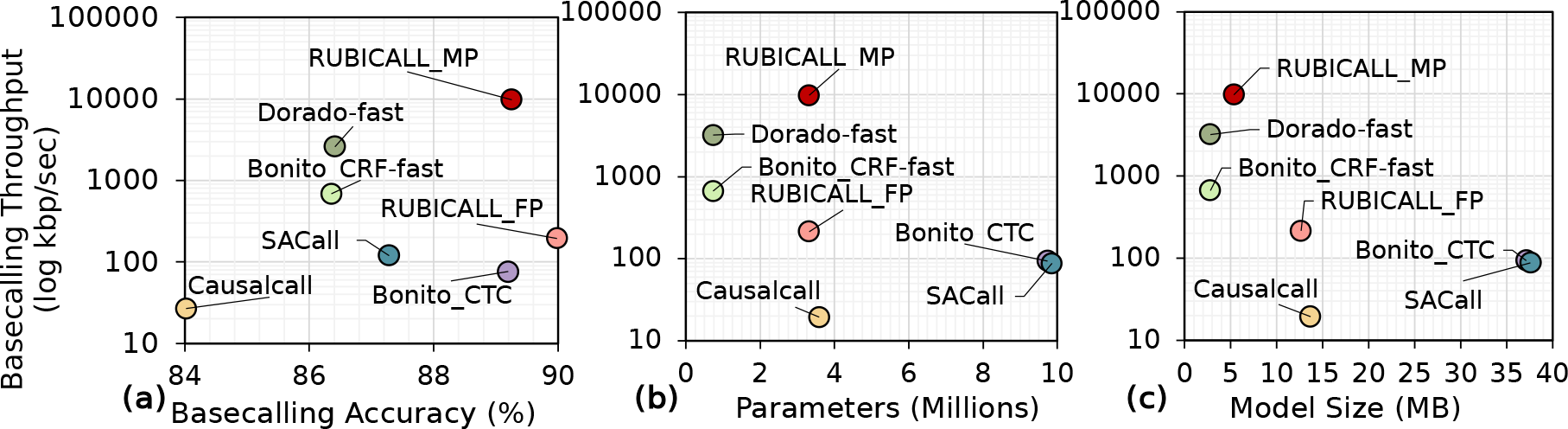
Comparison of average basecalling throughput for RUBICALL-MP with state-of-the-art basecallers in terms of: (a) average basecalling accuracy, (b) model parameters, and (c) model size. RUBICALL-MP provides higher compute performance with lower model size when compared to RUBICALL-FP because of the mixed-precision computation.

We make six key observations. First, compared to `Dorado-fast`, the fastest basecaller, `RUBICALL-MP` provides, on average, 2.85% higher accuracy with 3.77× higher basecalling throughput. Therefore, `RUBICALL-MP` provides both accuracy and high basecalling throughput. Second, `RUBICALL-MP` provides 128.13× higher basecalling throughput without any loss in accuracy compared to `Bonito_CTC`, which is an expert-designed basecaller. Unlike `Bonito_CTC`, This is because `RUBICALL-MP` has a mixed precision neural architecture that leads to high compute density. Third, by using mixed-precision quantization, `RUBICALL`-`MP` provides 50.15× higher performance when compared to its floating-point implementation (`RUBICALL-FP`). Fourth, `SACall` has the highest number of neural network model parameters, which are 2.74×, 13.49×, 1.01×, 13.49×, and 2.97× more than `Causalcall`, `Bonito_CRF-fast`, `Bonito_CTC`, `Dorado-fast`, and `RUBICALL-MP`, respectively. `SACall` uses a large transformer model with an attention mechanism that leads to an over-parameterized model. Fifth, `Dorado-fast` has 4.92×, 13.33×, 13.49×, and 4.54× lower number of trainable model parameters than `Causalcall`, `Bonito_CTC`, `SACall`, and `RUBICALL-MP`. As discussed earlier, `Dorado-fast` provides 2.85% lower accuracy with 3.77× lower basecalling throughput. While `Dorado-fast` has a 4.54× lower number of trainable model parameters, the difference in model size is only 1.92× because `RUBICALL-MP` has each layer quantized to a different precision. Sixth, compared to basecallers with skip connections, `RUBICALL-MP` provides 2.55× and 6.93× smaller model size compared to `Causalcall` and `Bonito_CTC`, respectively. The decrease in model size is due to: (1) a lower number of neural network layers; and (2) optimum bit-width precision for each neural network layer. Sixth, all the baseline basecallers use floating-point arithmetic precision for all neural network layers. This leads to very high memory bandwidth and processing demands. We conclude that `RUBICALL-MP` provides the ability to basecall quickly, and efficiently scale basecalling by providing reductions in both model size and neural network model parameters.

### 2.3. Performance Comparison

We compare the speed of `RUBICALL-MP` against baseline basecallers in Figure 6. We make three major observations. First, `RUBICALL-MP` consistently outperforms all the other basecallers for all the evaluated species. `RUBICALL-MP` improves average performance by 364.89×, 14.25×, 128.13×, 81.58×, and 3.77× over `Causalcall`, `Bonito_CRF-fast`, `Bonito_CTC`, `SACall`, and `Dorado-fast`, respectively. Second, as `RUBICALL`-`MP` each layer is quantized to a different precision, it provides 50.15× higher performance when compared to its floating-point only implementation (`RUBICALL-FP`). Third, `RUBICALL-FP`, by using floating-point precision, provides 7.28×, 2.56×, and 1.63× higher performance compared to `Causalcall`, `Bonito_CTC`, and `SACall`, respectively. Figure S7 in Supplementary Section S5 demonstrates the performance of all the evaluated basecallers on NVIDIA A40 [70] GPU. We conclude that using mixed-precision computation, `RUBICALL-MP` consistently performs better than the baseline basecallers.

**Figure 6:**
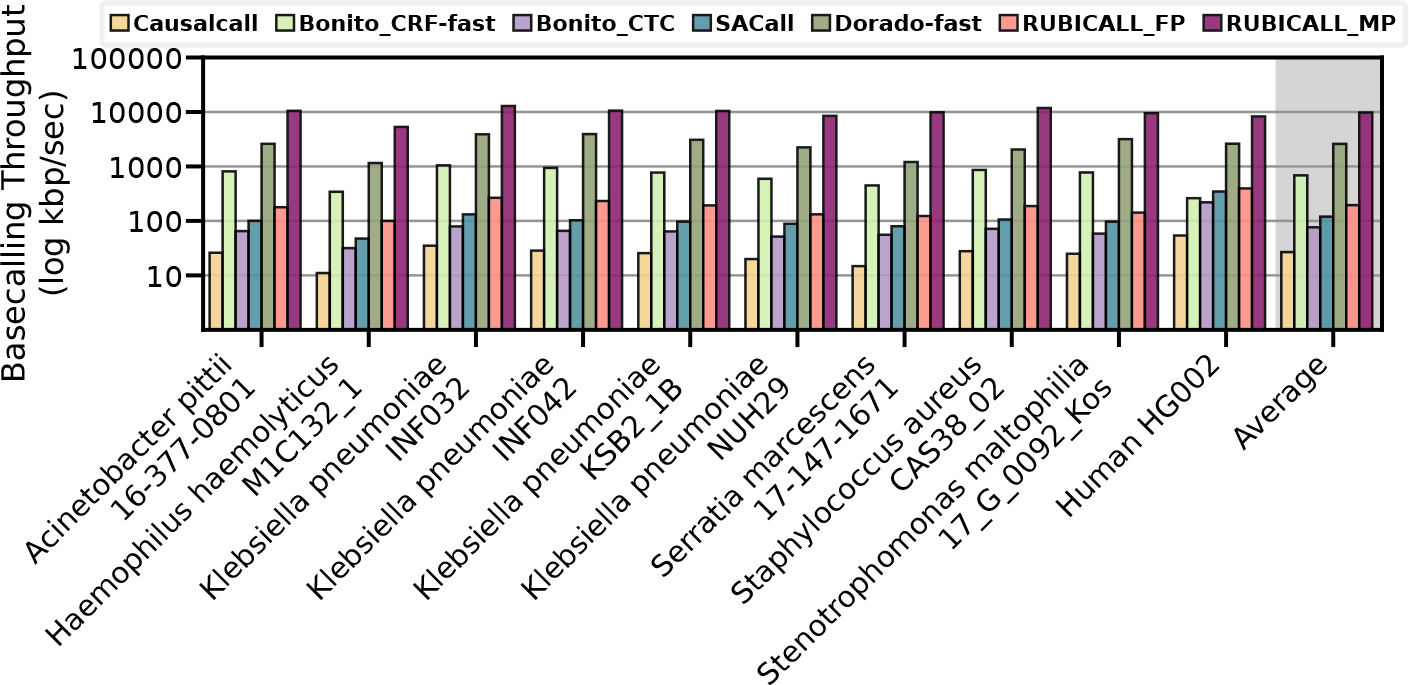
Performance comparison of RUBICALL (using floating-point precision (RUBICALL-FP) and mixed-precision (RUBICALL-MP)) and five state-of-the-art basecallers on AMD MI210. The y-axis is on a logarithmic scale.

### 2.4. Basecalling Accuracy

We compare the basecalling accuracy of `RUBICALL` against baseline basecallers in Figure 7. `RUBICALL-MP` and `RUBICALL-FP` use the same model architecture and produce the same basecalled reads, so we report results as `RUBICALL`. We make three major observations. First, compared to `Dorado-fast` and `Bonito_CRF-fast`, we observe `RUBICALL` achieves 2.85% and 2.89% higher accuracy over these RNN-based basecallers, respectively. `RUBICALL` provides 5.23% and 0.06% higher accuracy than CNN-based basecaller `Causalcall` and `Bonito_CTC`, respectively. Compared to a state-of-the-art transformer-based basecaller, `SACall`, `RUBICALL` achieves 1.97% higher basecalling accuracy. Second, `Bonito_CTC` has 2.93× higher parameters (Figure 5(a)) while having similar accuracy as `RUBICALL`. Third, `Causalcall` and `SACall` are unable to align half of *Haemophilus haemolyticus M1C132_1* reads to its reference. Therefore, it is deemed unaligned and cannot be used to determine its read accuracy. We conclude that `RUBICALL` provides the highest accuracy compared to other basecallers.

**Figure 7:**
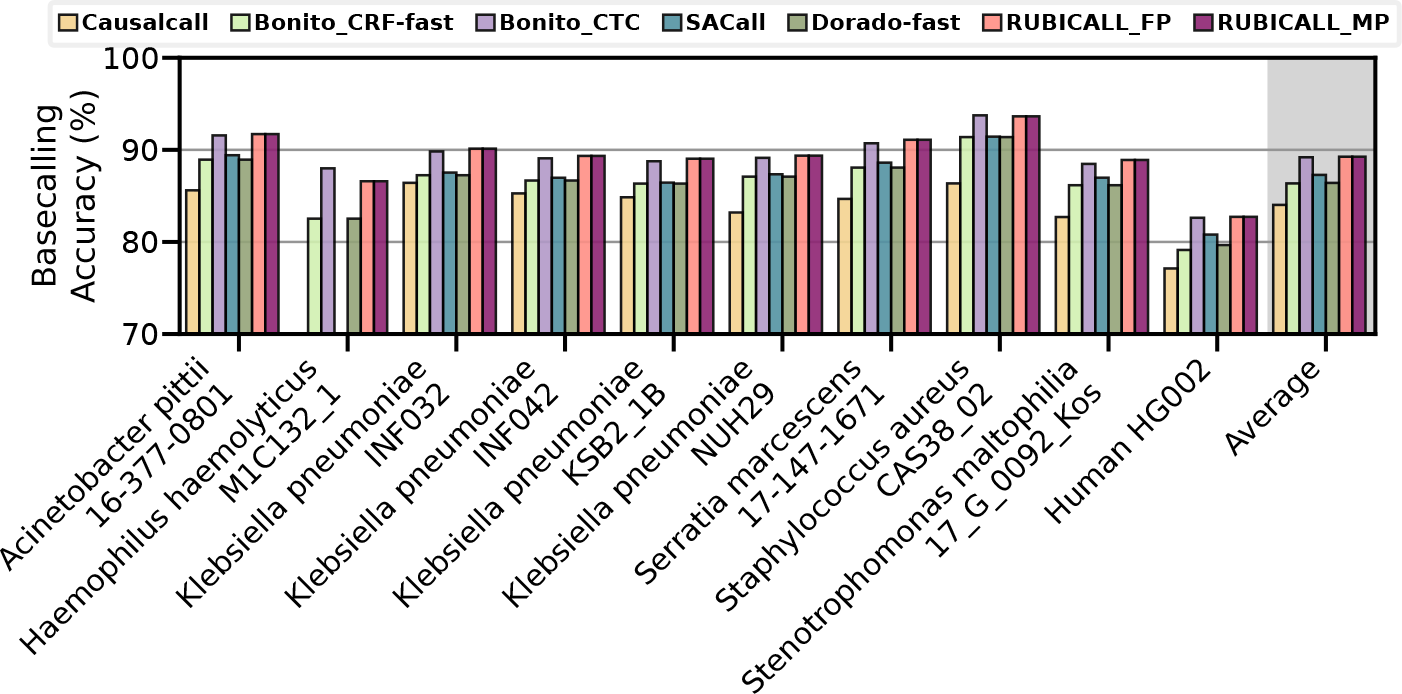
Basecalling accuracy comparison of RUBICALL (using floating-point precision (RUBICALL-FP) and mixed-precision (RUBICALL-MP)).

### 2.5. Downstream Analysis

#### 2.5.1. De Novo Assembly

We provide the statistics related to the accuracy, completeness, and contiguity of assemblies we generate using the basecalled reads from `Causalcall`, `Bonito_CRF-fast`, `Bonito_CTC`, `SACall`, `Dorado-fast`, and `RUBICALL` in Table 1. For `Genome Fraction (%)`, `Average Identity (%)`, and Quality Value (QV), we highlight the highest achieved value. While for `Assembly Length`, `Average GC (%)`, and `NG50`, we highlight the value closest to the real assembly length. For `Total Indels` and `Indel Ratio (%)`, the best-performing basecaller has the lowest value. We also collect the number of unique k-mers and the frequency of each unique k-mer in a given sequence to perform a comparison of under and over-represented k-mers in Supplementary Section S7.

**Table 1:** Assembly quality comparison of the evaluated basecallers for different species. We measure assembly accuracy in terms of genome fraction (Genome Fraction (%)) and average identity (Average Identity (%)). Genome fraction is the portion of the Reference genome that can align to a given assembly, while average identity is the average of the identity of assemblies when compared to their respective Reference genomes. We measure statistics related to the contiguity and completeness of the assemblies in terms of the overall assembly length (Assembly Length), Average GC content (Average GC (%)) (i.e., the ratio of G and C bases in an assembly), NG50 statistics (NG50) (i.e., shortest contig at the half of the overall Reference genome length), total number of indels in all aligned bases in the assembly (Total Indels), the ratio of indels to assembly length (Indel Ratio (%)), and the reliability of basepairs using the quality value (Quality Value). NA indicates that the generated assemblies were unalignable to the reference genome.

| Dataset | Basecaller | Genome Fraction (%) | Average Identity (%) | Assembly Length | Average GC (%) | NG50 | Total Indels | Indel Ratio (%) | Quality Value (QV) |
| --- | --- | --- | --- | --- | --- | --- | --- | --- | --- |
| Acinetobacter pittii 16-377-0801 | Causalcall | 92.45 | 86.18 | 3,826,077 | 42.23 | 3,826,077 | 270,228 | 7.06 | 11.99 |
|  | Bonito_CRF-fast | 96.64 | 89.29 | 3,628,317 | 38.82 | 3,628,317 | 242,373 | 6.68 | 12.03 |
|  | Bonito_CTC | 96.87 | 91.44 | 3,676,821 | 38.9 | 3,676,821 | 210,496 | 5.72 | 12.45 |
|  | SACall | 96.68 | 89.42 | 3,699,232 | 38.7 | 3,699,232 | 247,997 | 6.7 | 12.1 |
|  | Dorado-fast | 96.37 | 88.72 | 3,839,847 | 39.09 | 3,839,847 | 245,016 | 6.38 | 12.03 |
|  | RUBICALL | 96.87 | 91.51 | 3,694,086 | 38.82 | 3,694,086 | 208,748 | 5.65 | 15.42 |
|  | Reference | 100 | 100 | 3,814,719 | 38.78 | 3,814,719 | 0 | 0 | - |
| Haemophilus haemolyticus M1C132_1 | Causalcall | 0.00 | 0.00 | 0 | 0 | 0 | 0 | 0 | NA |
|  | Bonito_CRF-fast | 88.76 | 91.51 | 2,046,024 | 37.98 | 2,046,024 | 128,481 | 6.28 | 12.25 |
|  | Bonito_CTC | 96.87 | 90.70 | 1,957,480 | 38.87 | 1,957,480 | 118,253 | 6.04 | 15.34 |
|  | SACall | 90.11 | 88.45 | 2,032,994 | 38.22 | 1,880,730 | 134,702 | 6.63 | 13.15 |
|  | Dorado-fast | 89.42 | 88.97 | 2,110,860 | 39.49 | 2,110,860 | 129,503 | 6.14 | 12.38 |
|  | RUBICALL | 96.87 | 90.54 | 1,966,781 | 38.92 | 1,966,781 | 119,777 | 6.09 | 15.37 |
|  | Reference | 100 | 100 | 2,042,591 | 38.46 | 2,042,591 | 0 | 0 | - |
| Klebsiella pneumoniae INF032 | Causalcall | 92.45 | 87.35 | 4,959,127 | 56.9 | 4,959,127 | 353,550 | 7.13 | 10.54 |
|  | Bonito_CRF-fast | 92.69 | 87.53 | 4,761,297 | 57.19 | 4,761,297 | 347,299 | 7.29 | 10.56 |
|  | Bonito_CTC | 94.50 | 90.20 | 4,897,352 | 56.65 | 4,897,352 | 317,428 | 6.48 | 11.26 |
|  | SACall | 93.97 | 88.08 | 4,874,880 | 56.87 | 4,874,880 | 379,028 | 7.78 | 10.8 |
|  | Dorado-fast | 93.00 | 87.69 | 5,063,562 | 56.8 | 5,063,562 | 348,572 | 6.88 | 10.64 |
|  | RUBICALL | 94.51 | 90.30 | 4,924,240 | 56.85 | 4,924,240 | 314,651 | 6.39 | 11.27 |
|  | Reference | 100 | 100 | 5,111,537 | 57.63 | 5,111,537 | 0 | 0 | - |
| Klebsiella pneumoniae INF042 | Causalcall | 91.44 | 87.36 | 5,288,166 | 56.94 | 5,288,166 | 374,162 | 7.08 | 10.84 |
|  | Bonito_CRF-fast | 92.08 | 88.49 | 5,052,889 | 56.8 | 5,052,889 | 357,354 | 7.07 | 10.93 |
|  | Bonito_CTC | 93.12 | 90.49 | 5,111,083 | 56.61 | 5,111,083 | 317,075 | 6.2 | 11.40 |
|  | SACall | 92.93 | 88.60 | 5,149,039 | 56.72 | 5,149,039 | 369,388 | 7.17 | 11.08 |
|  | Dorado-fast | 90.21 | 88.20 | 5,737,059 | 56.44 | 5,401,717 | 342,141 | 5.96 | 10.98 |
|  | RUBICALL | 93.12 | 90.60 | 5,146,050 | 56.72 | 5,146,050 | 312,448 | 6.07 | 11.42 |
|  | Reference | 100 | 100 | 5,337,491 | 57.41 | 5,337,491 | 0 | 0 | - |
| Klebsiella pneumoniae KSB2_1B | Causalcall | 91.58 | 86.97 | 5,175,311 | 57.09 | 5,175,311 | 363,807 | 7.03 | 10.88 |
|  | Bonito_CRF-fast | 90.24 | 88.00 | 4,932,626 | 56.71 | 4,932,626 | 357,769 | 7.25 | 10.86 |
|  | Bonito_CTC | 93.07 | 90.11 | 5,003,377 | 56.69 | 5,003,377 | 320,519 | 6.41 | 11.41 |
|  | SACall | 93.58 | 88.19 | 5,034,408 | 56.79 | 5,034,408 | 372,380 | 7.4 | 11.16 |
|  | Dorado-fast | 90.28 | 87.67 | 5,442,186 | 56.72 | 5,261,731 | 349,387 | 6.42 | 11.03 |
|  | RUBICALL | 93.07 | 89.89 | 5,023,639 | 56.75 | 4,932,626 | 357,769 | 7.12 | 11.25 |
|  | Reference | 100 | 100 | 5,228,889 | 57.59 | 5,228,889 | 0 | 0 | - |
| Klebsiella pneumoniae NUH29 | Causalcall | 89.08 | 86.01 | 5,158,874 | 56.78 | 5,158,874 | 389,676 | 7.55 | 11.75 |
|  | Bonito_CRF-fast | 92.17 | 89.34 | 4,942,833 | 57.01 | 4,942,833 | 355,690 | 7.2 | 11.47 |
|  | Bonito_CTC | 94.36 | 90.26 | 4,918,147 | 57.04 | 4,918,147 | 324,406 | 6.6 | 11.92 |
|  | SACall | 93.66 | 88.58 | 4,978,307 | 57.06 | 4,978,307 | 360,950 | 7.25 | 11.56 |
|  | Dorado-fast | 92.27 | 88.12 | 5,195,594 | 57.01 | 5,195,594 | 355,728 | 6.85 | 11.56 |
|  | RUBICALL | 94.36 | 90.43 | 4,940,813 | 57.18 | 4,940,813 | 316,019 | 6.4 | 11.83 |
|  | Reference | 100 | 100 | 5,134,281 | 57.61 | 5,134,281 | 0 | 0 | - |
| Serratia marcescens 17-147-1671 | Causalcall | 89.91 | 86.23 | 5,532,953 | 57.86 | 5,422,052 | 401,545 | 7.26 | 13.39 |
|  | Bonito_CRF-fast | 96.06 | 89.56 | 5,479,812 | 58.85 | 5,282,474 | 345,351 | 6.3 | 12.66 |
|  | Bonito_CTC | 96.76 | 91.38 | 5,534,329 | 58.41 | 5,316,651 | 298,982 | 5.4 | 13 |
|  | SACall | 94.29 | 89.36 | 5,366,913 | 58.57 | 5,366,913 | 358,954 | 6.69 | 12.27 |
|  | Dorado-fast | 96.51 | 88.87 | 5,758,989 | 58.29 | 5,282,474 | 348,968 | 6.06 | 12.5 |
|  | RUBICALL | 96.76 | 91.59 | 5,597,251 | 58.52 | 5,346,640 | 294,643 | 5.26 | 13.01 |
|  | Reference | 100 | 100 | 5,517,578 | 59.13 | 5,517,578 | 0 | 0 | - |
| Staphylococcus aureus CAS38_02 | Causalcall | 94.35 | 87.29 | 2,849,123 | 36.59 | 2,810,038 | 191,730 | 6.73 | 10.8 |
|  | Bonito_CRF-fast | 96.27 | 91.49 | 2,790,895 | 33.05 | 2,752,169 | 149,623 | 5.36 | 11.59 |
|  | Bonito_CTC | 97.03 | 93.57 | 2,858,986 | 32.86 | 2,819,356 | 123,542 | 4.32 | 12.82 |
|  | SACall | 95.66 | 91.25 | 2,837,503 | 32.91 | 2,798,079 | 165,200 | 5.82 | 11.57 |
|  | Dorado-fast | 96.70 | 91.16 | 2,927,882 | 33.52 | 2,752,169 | 152,216 | 5.2 | 11.64 |
|  | RUBICALL | 97.03 | 93.36 | 2,860,885 | 33.24 | 2,821,276 | 124,795 | 4.36 | 12.59 |
|  | Reference | 100 | 100 | 2,902,076 | 32.82 | 2,902,076 | 0 | 0 | - |
| Stenotrophomonas maltophilia 17_G_0092_Kos | Causalcall | 94.85 | 85.73 | 4,823,177 | 63.66 | 4,823,177 | 366,228 | 7.59 | 11.01 |
|  | Bonito_CRF-fast | 94.60 | 89.74 | 4,596,898 | 65.5 | 4,596,898 | 337,040 | 7.33 | 11.10 |
|  | Bonito_CTC | 95.42 | 90.14 | 4,664,226 | 64.82 | 4,664,226 | 298,711 | 6.4 | 11.51 |
|  | SACall | 95.28 | 88.50 | 4,672,540 | 64.98 | 4,672,540 | 339,853 | 7.27 | 11.11 |
|  | Dorado-fast | 92.99 | 87.70 | 4,854,007 | 63.99 | 4,854,007 | 337,105 | 6.94 | 11.01 |
|  | RUBICALL | 95.46 | 90.49 | 4,693,744 | 65.03 | 4,693,744 | 289,073 | 6.16 | 11.63 |
|  | Reference | 100 | 100 | 4,802,733 | 66.28 | 4,802,733 | 0 | 0 | - |
| Human HG002 | Causalcall | NA | NA | 130,962 | 42.95 | 13,522 | NA | NA | NA |
|  | Bonito_CRF-fast | 0.002 | 92.36 | 119,570,537 | 40.34 | 368,848 | 2,860 | 0 | 18.87 |
|  | Bonito_CTC | 0.430 | 95.06 | 134,732,516 | 40.86 | 371,590 | 384,243 | 0.29 | 18.58 |
|  | SACall | NA | NA | 63,025,520 | 39.87 | 320,873 | NA | NA | NA |
|  | Dorado-fast | 0.001 | 93.15 | 121,146,376 | 39.8 | 361,677 | 926 | 0 | 17.46 |
|  | RUBICALL | 0.125 | 94.50 | 140,928,248 | 40.99 | 393,950 | 100,256 | 0.1 | 17.81 |
|  | Reference | 100 | 100 | 2,947,743,500 | 40.79 | 2,947,743,500 | 0 | 0 | - |

We make six key observations. First, assemblies constructed using reads basecalled by `RUBICALL` provide the best reference genome coverage for *all* datasets (“Genome Fraction” in Table 1). This means that assemblies built using `RUBICALL`-basecalled reads are more complete than assemblies built using reads from other basecallers since a larger portion of the corresponding reference genomes align to their assemblies using `RUBICALL`-basecalled reads compared to that of using reads from other basecallers. Second, assemblies constructed using the `RUBICALL` reads usually have a higher average identity than that of `Causalcall`, `Bonito_CRF-fast`, `Bonito_CTC`, `SACall`, and `Dorado-fast`. These average identity results are tightly in line with the basecalling accuracy results we show in Figure 7. Although `Bonito_CRF-fast` provides a higher average identity for the Haemophilus haemolyticus M1C132_1 dataset (i.e., 91.51%), the genome coverage provided by both `Bonito_CRF-fast` and `Dorado-fast` is 2.2% lower than that provided by `RUBICALL` for the same dataset. This means a large portion of the assembly provided by `Bonito_CRF-fast` has low-quality regions as the reference genome cannot align to these regions due to high dissimilarity. Third, assemblies constructed using the `RUBICALL` reads provide better completeness and contiguity as they have 1) assembly lengths closer to their corresponding reference genomes and 2) higher NG50 results in most cases than those constructed using the `Bonito_CRF-fast` and `Bonito_CTC` reads. Fourth, although `Causalcall` usually provides the best results in terms of the assembly lengths and NG50 results, we suspect that these high NG50 and assembly length results are caused due to highly repetitive and inaccurate regions in these assemblies due to their poor genome fraction and average GC content results. The average GC content of the assemblies constructed using the `Causalcall` reads is significantly distant from the GC content of their corresponding reference genomes in most cases. This poor genome fraction and average GC content results suggest that such large NG50 and assembly length values from `Causalcall` may also be caused by poorly basecalled reads that lead to unresolved repetitive regions (i.e., bubbles in genome assembly graphs) or a strong bias toward certain error types (i.e., homopolymer insertions of a certain base) in the assembly [71, 72]. Fifth, the low Total Indels and Indel Ratio (%) for `RUBICALL` in an assembled sequence signify a sequence that closely resembles the expected reference with minimal insertions and deletions (indels). This indicates a well-structured and high-quality assembly. Such assemblies offer a clear and accurate representation of the original sequence, facilitating downstream analyses, gene prediction, functional annotation, and comparative genomics. Sixth, `RUBICALL` consistently provides a higher quality value (QV), indicating a low probability of sequencing errors. Therefore, compared to the other evaluated basecallers, `RUBICALL` has higher reliability of the assembled genome.

We conclude that, in most cases, the reads basecalled by `RUBICALL` lead to higher quality, more contiguous, and more complete assemblies than that provided by other state-of-the-art basecallers, `Causalcall`, `Bonito_CRF-fast`, `Bonito_CTC`, `SACall`, and `Dorado-fast`.

#### 2.5.2. Read Mapping

We provide the comparison of `RUBICALL` with `Causalcall`, `Bonito_CRF-fast`, `Bonito_CTC`, `SACall`, and `Dorado-fast` in terms of the total number of base mismatches, the total number of mapped bases, the total number of mapped reads, and the total number of unmapped reads in Figure 8(a), 8(b), 8(c), and 8(d), respectively. We also show the average read length, the overall number of mapped reads and the mapped bases, and the ratio of the number of mapped bases to the number of mapped reads in Supplementary Table S2.

**Figure 8:**
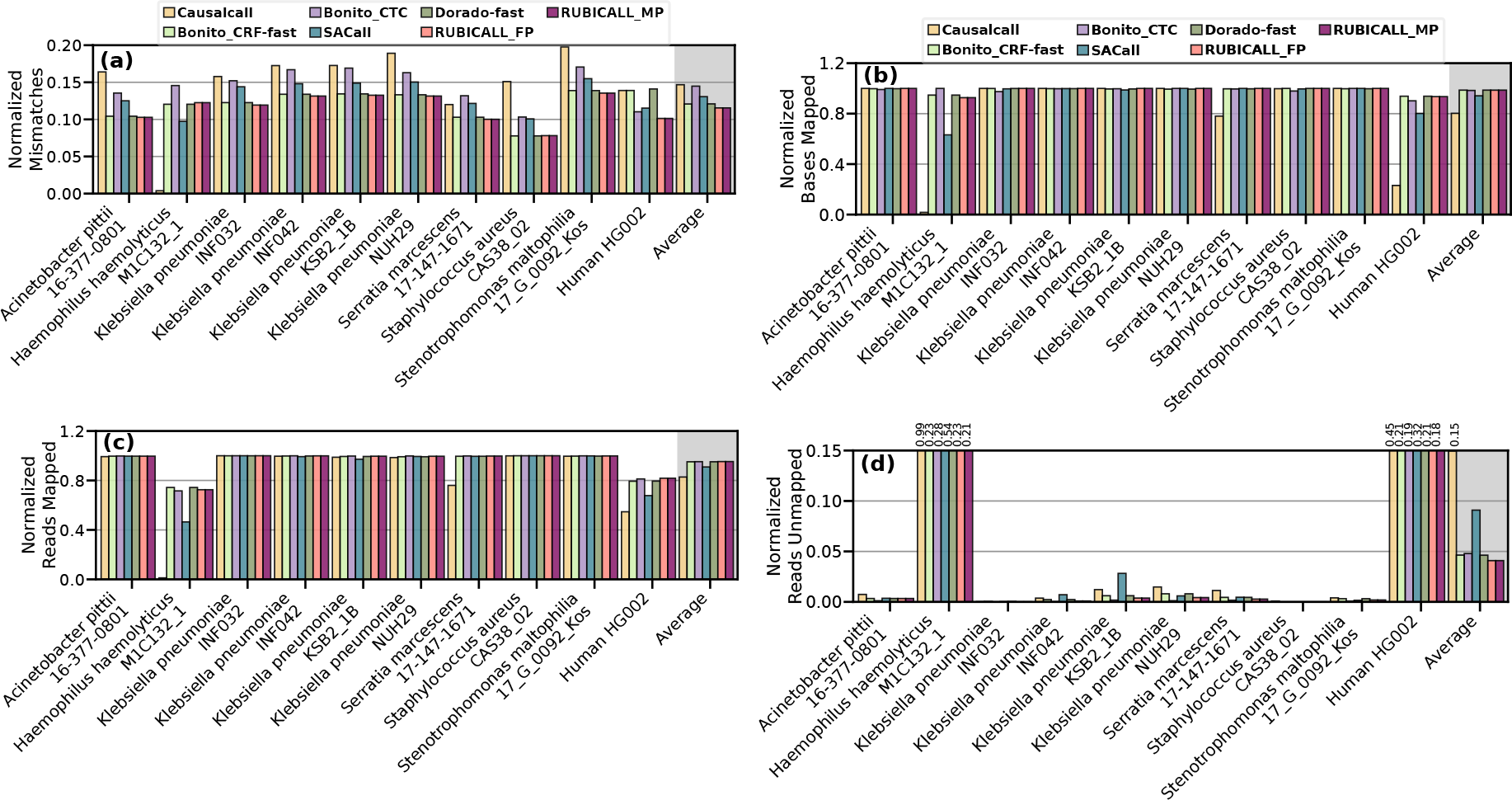
Comparison of RUBICALL (using floating-point precision (RUBICALL-FP) and mixed-precision (RUBICALL-MP)) for normalized (a) mismatches, (b) bases mapped, (c) reads mapped, and (d) reads unmapped.

We make five key observations. First, `RUBICALL` provides the lowest number of base mismatches, which are 26.97%, 22.66%, 11.45%, 12.35%, and 23.58% lower compared to `Causalcall`, `Bonito_CRF-fast`, `Bonito_CTC`, `SACall`, and `Dorado-fast`, respectively. This indicates that `RUBICALL` provides more accurate basecalled reads that share large similarity with the reference genome. This is in line with the fact that `RUBICALL` provides the highest basecalling accuracy, as we evaluate in Section 2.4. Second, `RUBICALL` provides, on average, 22.86%, 0.24%, and 4.77% higher number of mapped bases compared to `Causalcall`, `Bonito_CTC`, and `SACall`, respectively, and only 0.3% and 0.4% lower number of mapped bases when compared to `Bonito_CRF-fast` and `Dorado-fast`, respectively. Mapping more bases to the target reference genome confirms that the careful design and optimizations we perform when building `RUBICALL` have no negative effects on the basecalling accuracy. Third, unlike `Causalcall`, `RUBICALL`, `Bonito_CRF-fast`, `Bonito_CTC`, `SACall`, and `Dorado-fast`, all provide a high number of mapped reads. However, `RUBICALL` is the only basecaller that provides high-quality reads that have the highest number of base matches and the lowest number of base mismatches. Fourth, `RUBICALL` achieves 72.66%, 11.79%, 14.63%, 55.02%, and 11.61% lower unmapped reads compared to `Causalcall`, `Bonito_CRF-fast`, `Bonito_CTC`, `SACall`, and `Dorado-fast` respectively. This indicates that using `Causalcall`, `Bonito_CRF-fast`, `Bonito_CTC`, `SACall`, and `Dorado-fast` wastes a valuable, expensive resource, i.e., sequencing data, by not mapping reads to the reference genome due to basecalling inaccuracies during basecalling. If a read is flagged as unmapped during read mapping, then this read is excluded from all the following analysis steps affecting the overall downstream analysis results. Fifth, for each dataset, we find that the ratio of the number of mapped bases to the number of mapped reads and the average length of the reads are mainly similar across all basecallers (Supplementary Table S2), while `Causalcall` has a substantially lower ratio for the human genome. This mainly indicates that unaligned bases across basecallers are mainly shared within the mapped reads, resulting in a similar number of mapped reads with similar average lengths as well as the ratio. We conclude that `RUBICALL` reads provides the highest-quality read mapping results with the largest number of mapped bases and mapped reads.

### 2.6. SkipClip Analysis

Figure 9 shows the effect of `SkipClip` on validation accuracy using three different strides at which we remove a skip connection from a block, i.e., the epoch interval at which `SkipClip` removes a skip connection from a block. We use our `QABAS`-designed model that has five blocks of skip connections. We highlight the number of epochs needed to remove all the skip connections for different strides. For example, `Stride 1` requires five epochs to remove all the skip connections, while `Stride 3` requires fifteen epochs. We make three observations. First, `Stride 1` converges faster to the baseline accuracy compared to `Stride 2` and `Stride 3`. By using `Stride 1`, we quickly remove all the skip connections (in five epochs) giving enough fine-tuning iterations for the model to recover its loss in accuracy. Second, all the strides show the maximum drop in accuracy (1.27%-2.88%) when removing skip connections from block 1 and block 4. We observe these blocks consist of the highest number of neural network model parameters due to the skip connections (30.73% and 25.62% of the total model parameters are present in skip connections in block 1 and block 4, respectively). Therefore, the model requires more training epochs to recover its accuracy after the removal of skip connections from these blocks. Third, a lower stride can get rid of skip connections faster than using a higher stride. However, all strides eventually converge to the baseline accuracy at the expense of more training iterations. We conclude that `SkipClip` provides an efficient mechanism to remove hardware-unfriendly skip connections without any loss in basecalling accuracy.

**Figure 9:**
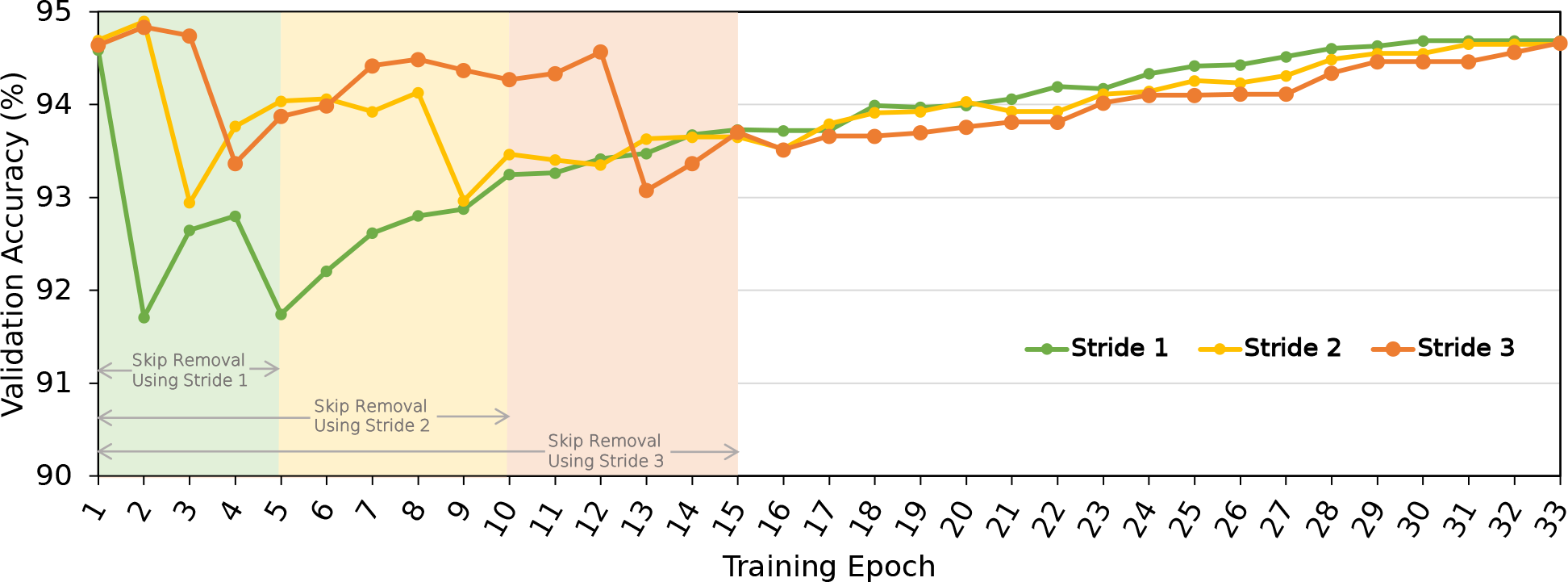
Effect of different strides while removing skip connections.

### 2.7. Effect of Pruning RUBICALL

Figure 10 shows the effect of pruning `RUBICALL` using two different pruning methods: unstructured element pruning and structured channel pruning.

**Figure 10:**
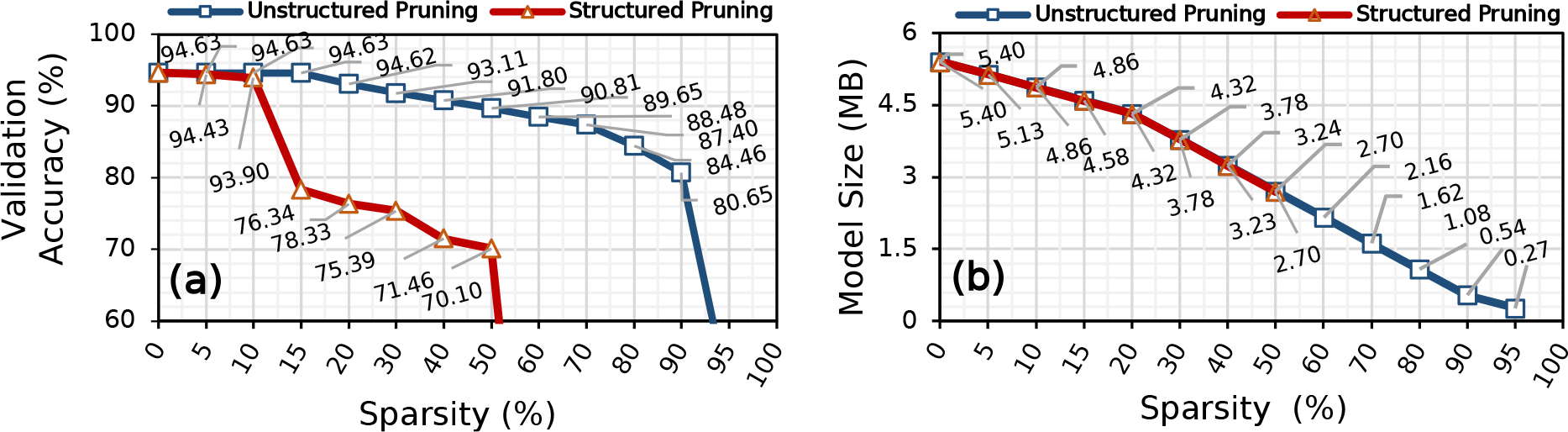
Effect of pruning RUBICALL on: (a) validation accuracy and (b) model size.

We make four major observations. First, we can remove up to 15% and 5% of model parameters providing 1.18% and 1.05% reductions in model size without any loss in accuracy by using unstructured pruning and structured pruning, respectively. However, unstructured pruning is unsuitable for hardware acceleration due to irregular structure, and structured pruning provides minimal model size (or parameters) savings. Therefore, we do not apply these pruning techniques to optimize `RUBICALL` further. Second, we observe a drop in accuracy for pruning levels greater than 15% and 5% for unstructured and structured pruning, respectively. This shows that `QABAS` found an optimal architecture as there is little room for pruning `RUBICALL` further without loss in accuracy.

Third, we observe that the *knee point* for unstructured pruning and structured pruning lies at 90% and 50%, where we achieve 80.65% and 70.10% of accuracy with 9.99× and 1.99× savings model size, respectively. After the knee point, we observe a sharp decline in accuracy. Fourth, below the knee point, we can trade accuracy for speed to further accelerate `RUBICALL` for hardware computation and resources by removing unimportant network weights. We conclude that pruning provides a tradeoff between accuracy and model size that can lead to further reductions in processing and memory demands for `RUBICALL`, depending on the type of device on which genomic analyses would be performed.

### 2.8. Explainability Into QABAS Results

We perform an explainability analysis to understand our results further and explain `QABAS`’s decisions. The search performed by `QABAS` provides insight into whether `QABAS` has learned meaningful representations in basecalling. In Figure 11(a) and 11(b), we extract the number of model parameters and precision of each parameter in a neural network layer to calculate the total size for each layer for `Bonito_CTC` and `RUBICALL-MP`, respectively. We highlight each layer’s precision (i.e., weights and activation precision) using distinct colors. Our range includes floating-point (i.e., `fp32`) computation to integer computation (i.e., `int16`, `int8`, and `int4`) for weight and activation. Based on our experiments in Section 2.1.2, we restrict the precision of weight and activation in `RUBICALL-MP` architecture in `QABAS` to `int8` and `int4`, respectively. We compare `RUBICALL-MP` to `Bonito_CTC` as it has the same backend (i.e, Quartznet [50]) and is designed by ONT experts. We make three observations. First, `QABAS` uses more bits in the initial layers than the final layers in `RUBICALL-MP`. `QABAS` learns that the input to `RUBICALL` uses an analog squiggle that requires higher precision, while the output is only the nucleotide bases (A, C, G, T), which can be represented using lower precision.

**Figure 11:**
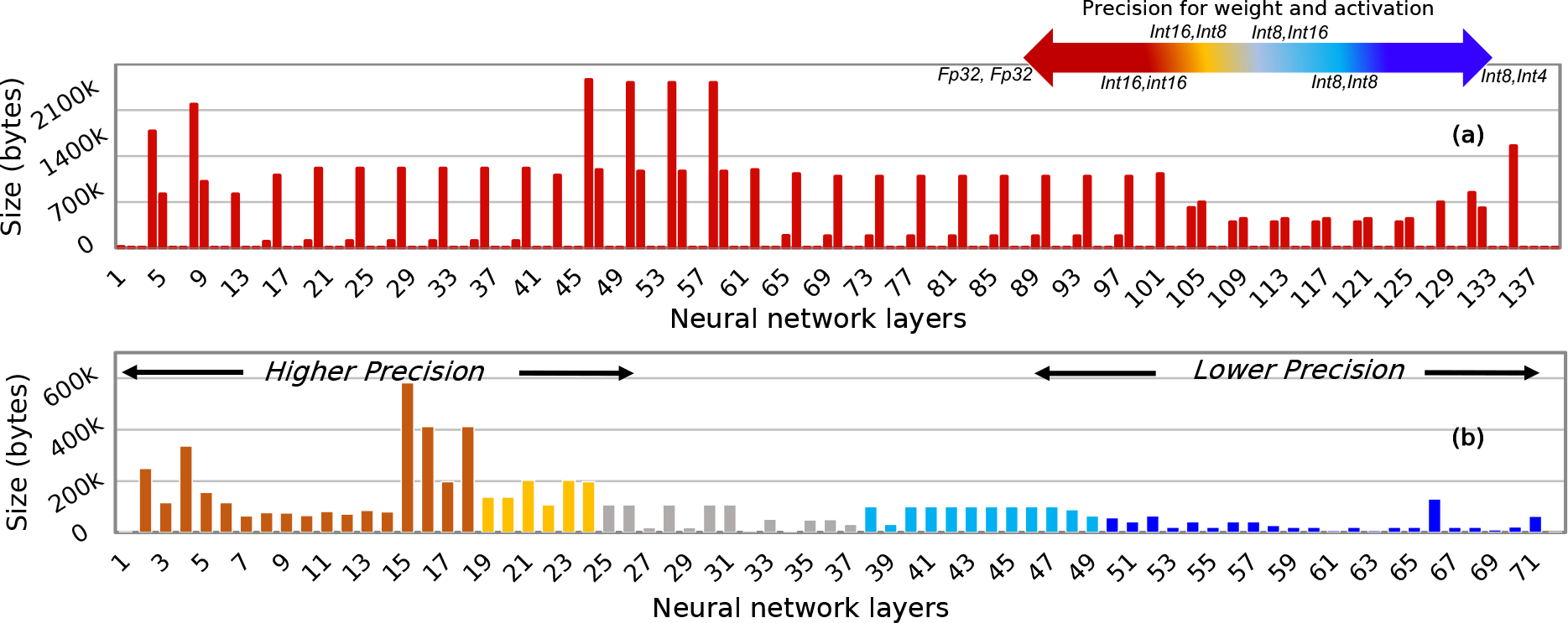
Layer size comparison for basecallers: (a) Bonito_CTC, and (b) RUBICALL-MP.

Second, `RUBICALL` uses 1.97× less number of neural network layers than `Bonito_CTC` while providing similar or higher basecalling accuracy on the evaluated species (Section 2.4). Thus, the superior performance of a basecaller architecture is not explicitly linked to its model complexity, and `QABAS`-designed models are parameter efficient. Third, `Bonito_CTC` uses the same single-precision floating-point representation (FP32) for all neural network layers, which leads to very high memory bandwidth and processing demands. Whereas `RUBICALL` has every layer quantized to a different quantization domain. We conclude that `QABAS` provides an efficient automated method for designing more efficient and hardware-friendly genomic basecallers compared to expert-designed basecallers.

## 3 Discussion

We are witnessing a tremendous transformation in high-throughput sequencing to significantly advance omics and other life sciences. The bioinformatics community has developed a multitude of software tools to leverage increasingly large and complex sequencing datasets. Deep learning models have been especially powerful in modeling basecalling.

### Importance of basecalling

Basecalling is the most fundamental computational step in the high-throughput sequencing pipeline. It is a critical problem in the field of genomics, and it has a significant impact on downstream analyses, such as variant calling and genome assembly. Improving the efficiency of basecalling has the potential to reduce the cost and time required for genomic analyses, which has practical implications for real-world applications. `RUBICALL` offers a valuable alternative for researchers and practitioners who seek a balance between accuracy and speed. By maintaining competitive accuracy levels while significantly improving speed, our framework addresses the needs of various applications with stringent time constraints, ultimately benefiting a broader range of users. We believe that RUBICON provides a significant improvement over existing methods, and it has practical implications for the genomics community.

### Need to improve the throughput of basecallers

Increasing throughput and reducing model size is critical because of the following three reasons. First, current basecallers already have high accuracy, but biologists do not pay attention to the throughput implications of using large deep learning-based models [31]. We observe researchers building larger and larger basecallers in an attempt to gain more accuracy without heeding to the disproportionately higher amount of power these basecallers are consuming. Moreover, none of the previous basecallers [29, 30, 40–43, 45, 73] have been optimized for mixed-precision execution to reduce energy consumption. As energy usage is proportional to the size of the network, energy-efficient basecalling is essential to enable the adoption of more and more sophisticated basecallers. Second, speed is critical in certain applications and use cases, particularly those that require real-time or near-real-time processing. `RUBICON` addresses these needs by focusing on hardware optimization and efficient implementation, ultimately enabling faster basecalling and potentially opening up new possibilities for applications with stringent time constraints. Third, as deep learning techniques and hardware continue to evolve, the balance between accuracy and speed/energy will remain an important aspect of model development. `RUBICON` provides a foundation for future research and innovation in hardware-friendly deep learning models for genomic basecalling.

### Evaluating RUBICON on other platforms

All the state-of-the-art basecallers and RUBICON use high-level libraries, such as PyTorch or TensorFlow, which abstract the hardware architecture and provide a unified interface for deep learning computations. These libraries work out-of-the-box for AMD GPUs and are equally optimized for them. Currently, high-level libraries do not provide capabilities to exploit low-precision tensor cores available on the latest GPUs. As a result, existing basecallers take advantage of comparable architectural capabilities regardless of the specific GPU employed. Therefore, the hardware and software optimizations are at the same level for all supported GPU-based platforms.

### Automating basecaller generation process

Modern basecallers generally employ convolution neural networks to extract features from raw genomic sequences. However, designing a basecaller comes with a cost that a neural network model can have many different computational elements making the neural network tuning a major problem. At present, the vast majority of deep learning-based basecallers are manually tuned by computational biologists through manual trial and error, which is time-consuming. To a large extent, basecallers are being designed to provide higher accuracy without considering the compute demands of such networks. Such an approach leads to computationally complex basecallers that impose a substantial barrier to performing end-to-end time-sensitive genomic analyses. This vast dependence of computational biologists and biomedical researchers on these deep learning-based models creates a critical need to find efficient basecalling architectures optimized for performance.

During our evaluation, we ran `QABAS` for 96 GPU hours to sample architectures from our search space. Using complete sampling to evaluate all the 1.8 10^32^ viable options would take at least ∼4.3 10^33^ GPU hours. Thus, `QABAS` accelerates the basecaller architecture search to develop high-performance basecalling architectures. The final model architecture can be further fine-tuned for other hyperparameters [74,75], such as learning rate and batch size (for example, with grid search or neural architecture search). Throughout our experiments, we build general-purpose basecalling models by training and testing the model using an official, open-source ONT dataset that consists of a mix of different species. We did not specialize basecalling models for a specific specie. Past works, such as [29], show that higher basecalling accuracy can be achieved by building species-specific models.

### Extending RUBICON

`RUBICON`’s modular design allows for the incorporation of additional layers or techniques, such as RNN, LSTM, and Transformers, to potentially increase accuracy further. We focus on convolution-based networks because: (a) matrix multiplication is the fundamental operation in such networks that is easily amenable to hardware acceleration, (b) the training and inference of RNN and LSTM models inherently involve sequential computation tasks, which poses a challenge for their acceleration on contemporary hardware such as GPUs and field-programmable gate arrays (FPGAs) [76–79], and (c) transformer-based models are typically composed of multiple fully connected layers, which can be supported in `RUBICON` by modifying convolutional layers for improved computational efficiency and performance [80]. As future work, `QABAS` can be extended in two ways: (1) evaluate advance model architectures (such as RNN, transformer, etc.), and (2) perform more fine-grain quantization. First, extending `QABAS` to other model architectures is important for researchers to quickly evaluate different computational elements. As the field of machine learning is rapidly evolving, it is non-trivial for researchers to adapt their models with the latest deep learning techniques. Second, currently, we perform mixed precision quantization, where every layer is quantized to a different domain. In the future, we can quantize every dimension of the weights to different precision. Such an approach would increase the design space of neural network architectural options to many folds. `QABAS` enables easy integration to explore such options automatically. Thus, `QABAS` is easily extensible and alleviates the designer’s burden in exploring and finding sophisticated basecallers for different hardware configurations. We would explore two future directions for pruning a basecaller. First, currently, we perform one-shot pruning, whereby we prune the model once and then fine-tune the model until convergence. Another approach could be to perform iterative pruning, where after every training epoch, we can re-prune the model using certain pruning criteria. Such an approach would further evaluate the fine-grained pruning limit of a basecaller. Second, an interesting future direction would be to combine multiple pruning techniques, e.g., structured channel pruning with structured group pruning (where we maintain the structure of the tensors without causing sparsity). Such an approach could lead to higher pruning ratios without substantial accuracy loss.

### Importance of RUBICALL beyond basecalling

For `SkipClip`, we demonstrate its applicability on basecalling only, while there are other genome sequencing tasks where deep learning models with skip connections are actively being developed, such as predicting the effect of genetic variations [73, 81], detecting replication dynamics [82], and predicting super-enhancers [83]. In Supplementary S2.1, we show the effect of manual skip removal, where we manually remove all the skip connections at once. We observe that the basecaller achieves 90.55% accuracy (4.08% lower than the baseline model with skip connections). By manual skip removal, the basecaller is unable to recover the loss in accuracy because CNN-based basecallers are sensitive to skip connections. Therefore, `SkipClip` provides a mechanism to develop hardware-friendly deep learning models for other genomic tasks.

### Separation between QABAS and SkipClip

Both `QABAS` and `SkipClip` share the overarching objective of creating a compact basecalling network without compromising accuracy. However, they approach this goal from distinct perspectives and employ different optimization tools. The following three points justify the separation of the two methods. First, skip connections are integral to stable model training, and by retaining them during the initial `QABAS` phase, we ensure effective training of the final basecalling network. The subsequent application of `SkipClip` allows for the controlled removal of skip connections, contributing to a more robust solution. Second, `QABAS` might find an architecture with skip connections, whereas `SkipClip` employs knowledge distillation for skip connection removal, addressing a specific aspect not efficiently handled by `QABAS` alone. Third, unlike `SkipClip`, `QABAS` tailors the neural network architecture for hardware efficiency without relying on a teacher network. The teacher network provides an upper bound on the achievable accuracy. Therefore, this two-step approach optimally combines the strengths of NAS and knowledge distillation, ensuring a comprehensive and effective optimization process for a compact and efficient basecalling model.

## 4 Conclusion

Nanopore sequencing generates noisy electrical signals that require conversion into a standard DNA nucleotide base string through a computational process known as basecalling. Efficient basecalling is crucial for subsequent genome analysis steps. Current basecalling approaches often neglect computational efficiency, resulting in slow, inefficient, and resource-intensive basecallers. To address this, we present `RUBICON`, a framework designed for creating hardware-optimized basecallers. `RUBICON` introduces two novel machine-learning techniques: `QABAS`, an automatic architecture search for computation blocks and optimal bit-width precision, and `SkipClip`, a dynamic skip connection removal module that significantly reduces resource and storage requirements without sacrificing basecalling accuracy. We demonstrate the capabilities of `QABAS` and `SkipClip` by designing `RUBICALL`, the first hardware-optimized basecaller, demonstrates fast, accurate, and efficient basecalling, achieving ∼ 6.88 × reductions in model size with 2.94 × fewer neural network parameters compared to an expert designed basecaller. We believe our open-source implementations of `RUBICON` will inspire advancements in genomics and omics research and development.

## 5 Methods

### Evaluation setup

Table 2 provides our system details. We evaluate `RUBICALL` using: (1) AMD MI210 GPU [69] (`RUBICALL-FP`) using floating-point precision computation, and (2) Versal ACAP VC2802 [65], a cutting-edge spatial vector computing system from AMD-Xilinx (`RUBICALL-MP`) using mixed-precision computation. The Versal ACAP VC2802 features Versal AI Engine ML (AIE-ML) [65] with 304 cores. The AIE-ML vector datapath implements two-dimensional single instruction, multiple data (SIMD) [84] operations using precisions ranging from int4×int8 to int16 × int16 operands that can execute 512 to 64 multiply-accumulate operations (MACs) per cycle, respectively. With its many different datatype precision options, AIE-ML acts as a suitable platform to demonstrate the benefits of a mixed precision basecaller. We train all the basecallers (`Causalcall`, `Bonito_CRF-fast`, `Bonito_CTC`, `SACall`, and `Dorado-fast`) using the same MI50 GPU. We use ONNX (Open Neural Network Exchange) [85] representation to evaluate the performance on AIE-ML by calculating bit operations (BOPs) [86], which measures the number of bitwise operations in a given network, take into account the total number of supported operations per datatype on AIE-ML.

**Table 2:** System parameters and hardware configuration for the CPU, GPU, and the AMD-Xilinx Versal ACAP.

|  |  |
| --- | --- |
| **CPU** | AMD EPYC 7742 [87]<br>@2.25GHz, 4-way SMT [88] |
| **Cache-Hierarchy** | 32×32 KiB L1-I/D, 512 KiB L2, 256 MiB L3 |
| **System Memory** | 4×32GiB RDIMM DDR4 2666 MHz [89] PCIe 4.0 ×128 |
| **OS details** | Ubuntu 21.04 Hirsute Hippo [90],<br>GNU Compiler Collection (GCC) version 10.3.0 [91] |
| **GPU** | AMD Radeon Instinct™ MI210 [69] 6656 Stream Processors@1.7GHz<br>64GB HBM2 PCIe 4.0 ×16, ROCm version 5.1.1 [92]<br>NVIDIA A40 [70] 10,752 CUDA Cores@1.2GHz, 48GiB DRAM<br>NVIDIA System Management Interface (NVIDIA-SMI) version 510.47.03 [93]<br>NVIDIA CUDA Compiler Driver (NVCC) version 11.4 [94] |
| **AMD-Xilinx Versal ACAP** | Versal ACAP VC2802 [65], 304×AIE-ML@1GHz,<br>19MB local memory, Dual-Core Arm Cortex-A72 [95] |

### QABAS setup details

We use the publicly available ONT dataset [63] sequenced using MinION Flow Cell (R9.4.1) for the training and validation during the `QABAS` search phase. The dataset comprises 1,221,470 reads, all sequenced from complete genomes. This ONT training dataset has an approximate list of 496 unique taxonomic IDs using the Kraken2 [96] taxonomic classification system [97]. We randomly select 30k samples from the training set for the search phase (specified using the `--chunks` parameter). We use nni [98] with nn-meter [99] to implement hardware-aware NAS. We use the Brevitas library [100] to perform quantization-aware training. The architectural parameters and network weights are updated using AdamW [101] optimizer with a learning rate of 2*e*^−3^, a beta value of 0.999, a weight decay of 0.01, and an epsilon of 1*e*^−8^. We set the hyperparameter *λ* to 0.6. We choose these values based on our empirical analysis. After the `QABAS` search phase, the sampled networks are trained until convergence with knowledge distillation using the same ONT dataset that we use during the `QABAS` search phase, with a batch size of 64, based on the maximum memory capacity of our evaluated Mi50 GPU. We set knowledge distillation hyperparameters alpha (*α*) and temperature (*τ*) at 0.9 and 2, respectively.

### QABAS search space

For the computations operations, we search for a design with one-dimensional (1D) convolution with ten different options: kernel size (KS) options (3, 5, 7, 9, 25, 31, 55, 75, 115, and 123) for grouped 1-D convolutions. We also use an identity operator that, in effect, removes a layer to get a shallower network. For quantization bits, we use bit-widths that are a factor of 2^*n*^, where 2<n<4 (since we need at least 2 bits to represent nucleotides A, C, G, T and 1 additional bit to represent an undefined character in case of a misprediction). We use four different quantization options for weights and activations (*<*`8,4`*>, <*`8,8`*>, <*`16,8`*>*, and *<*`16,16`*>*). We choose these quantization levels based on the precision support provided by our evaluated hardware and the effect of quantization on basecalling (Section 3). We use five different channel sizes with four repeats each. We choose the number of repeats based on the maximum memory capacity of our evaluated GPU. In total, we have ∼1.8×10^32^ distinct model options in our search space ℳ.

### SkipClip details

We use `Bonito CTC` as the teacher network, while the `QABAS`-designed model is the student network. We remove skip connections with a stride 1 (using parameter `--skip_stride`). Based on hyper-parameter tuning experiments (Supplementary S2.2), set knowledge distillation hyperparameters alpha (*α*) and temperature (*τ*) at 0.9 and 2, respectively. We use Kullback-Leibler divergence loss to calculate the loss [102].

### Pruning details

We use PyTorch [103] modules for both unstructured and structured pruning [104] with L1-norm, i.e., prune the weights that have the smallest absolute values. We apply one-shot pruning, where we first prune a model with a specific amount of sparsity, then train the model until convergence on the full ONT dataset [63].

### Baseline basecallers

`RUBICALL` is a pure convolution-based network. We focus on convolution-based networks because: (a) matrix multiplication is the fundamental operation in such networks that is easily amenable to hardware acceleration, (b) the training and inference of RNN and LSTM models inherently involve sequential computation tasks, which poses a challenge for their acceleration on contemporary hardware such as GPUs and field-programmable gate arrays (FPGAs) [76], and (c) transformer-based models are typically composed of multiple fully connected layers, which can be supported in `RUBICON` by modifying convolutional layers for improved computational efficiency and performance [80]. We compare `RUBICALL` against five different basecallers: (1) `Causalcall` [39] is a state-of-the-art basecaller with skip connections, (2) `Bonito_CRF-fast` [63] v0.6.2 is a recurrent neural network (RNN)-based version of basecaller from ONT that is optimized for throughput for real-time basecalling on Nanopore devices, (3) `Bonito_CTC` [63] v0.6.2 is convolutional neural network (CNN)-based hand-tuned basecaller from ONT, (4) `SACall` [44] is a transformer-based basecaller that uses an attention mechanism for basecalling, and (5) `Dorado-fast` [105] v0.4.0 is a LibTorch [106] version of `Bonito_CRF-fast` from ONT. `Dorado-fast` uses the same model architecture as `Bonito_CRF-fast` and uses the `Bonito` framework for model training. `Causalcall` and `Bonito_CTC` uses the same backend structure as `RUBICALL` (i.e., Quartznet [50]). We are aware of other basecallers such as `Halcyon` [43], `Helix` [41], and `Fast-bonito` [42]. However, these basecallers are either not open-source or do not provide training code with support for specific read formats.

### Basecalling reads

To evaluate basecalling performance, we use a set of reads generated using a MinION R9.4.1 flowcell. We use only R9 chemistry datasets as, currently, ONT does not provide a suitable public training dataset for R10 chemistry. They offer in-house trained R10 models that cannot be employed for a consistent evaluation across all basecallers. R9 and R10 chemistries involve distinct generations of nanopore technologies, including different pore proteins and read lengths. Therefore, models trained on R9 chemistry are incompatible for inference on R10 sequenced datasets. Due to these technical constraints, our study is currently limited to utilizing the available R9 chemistry training dataset from ONT and conducting inference exclusively on R9 chemistry datasets. Table 3 provides details on different organisms used in our evaluation. We use several bacterial species and the human genome. For Human HG002, we use 3 × depth of coverage.

**Table 3:** Details of datasets used in evaluation.

| Organism | Chemistry | # Reads | Reference Genome Size (bp) |
| --- | --- | --- | --- |
| Acinetobacter pittii 16-377-0801 | R9.4.1 | 4,467 | 3,814,719 |
| Haemophilus haemolyticus M1C132_1 | R9.4 | 8,669 | 2,042,591 |
| Klebsiella pneumoniae INF032 | R9.4 | 15,154 | 5,111,537 |
| Klebsiella pneumoniae INF042 | R9.4 | 11,278 | 5,337,491 |
| Klebsiella pneumoniae KSB2_1B | R9.4 | 15,178 | 5,228,889 |
| Klebsiella pneumoniae NUH29 | R9.4 | 11,047 | 5,134,281 |
| Serratia marcescens 17-147-1671 | R9.4.1 | 16,847 | 5,517,578 |
| Staphylococcus aureus CAS38_02 | R9.4.1 | 16,742 | 2,902,076 |
| Stenotrophomonas maltophilia 17_G_0092_Kos | R9.4 | 16,010 | 4,802,733 |
| Human HG002 | R9.4.1 | 300,000 | 2,947,743,500 |

Prior to basecalling, raw nanopore signals undergo a preprocessing pipeline to prepare them for input into the neural network. Raw nanopore signals, which can be hundreds of thousands of data points long, are normalized to ensure consistent input characteristics for the subsequent processing steps. We use empirically determined normalization scaling factors from ONT’s `Bonito_CTC` basecaller. The normalized signals are chunked into smaller segments, typically with overlapping regions. The chunk size and overlap are empirically set to 4000 bps and 500, respectively. Chunk size affects the balance between processing speed and accuracy. Smaller chunk sizes can lead to more accurate basecalling but may require more computational resources and time. Larger chunk sizes may be faster but can potentially introduce errors if the signal varies significantly within the chunk. Overlap represents the degree to which consecutive chunks share data with each other. Overlapping chunks can help mitigate the potential issues caused by abrupt changes in the signal at chunk boundaries. It allows for a smoother transition between chunks, reducing the chances of missing important information in the signal. However, a larger overlap may increase computational demands and processing time. After basecalling, the basecalled sequences obtained from individual signal segments are stitched back together to reconstruct the entire nucleotide sequence. The stitched sequences are then decoded to obtain the final basecalled sequences. We use the beam-search decoding [107] method to obtain the final basecalled sequences from stitched segments.

### Basecaller evaluation metrics

We evaluate the performance of `RUBICALL` using two different metrics: (1) basecalling throughput (kbp/sec), i.e., the throughput of a basecaller in terms of kilo basepairs generated per second, and (2) basecalling accuracy (%), i.e., the total number of bases of a read that are exactly matched to the bases of the reference genome divided by the total length of its alignment including insertions and deletions. We measure the basecalling throughput for the end-to-end basecalling calculations, including reading FAST5 files and writing out FASTQ or FASTA file using Linux */usr/bin/time -v* command. For basecalling accuracy, we align each basecalled read to its corresponding reference genome of the same species using the state-of-the-art read mapper, minimap2 [108]. We use Rebaler [109] to generate a consensus sequence from each basecalled read set, which replaces portions of the reference genome with read-derived sequences. The assembled genome is then polished with multiple rounds of Racon [110]. This results in an assembled genome that accurately represents the original data while minimizing potential errors introduced by the reference.

### Downstream analysis

We evaluate the effect of using `RUBICALL` and other baseline basecallers on two widely-used downstream analyses, *de novo* assembly [111] and read mapping [112].

#### *De novo* assembly

We construct *de novo* assemblies from the basecalled reads and calculate the statistics related to the accuracy, completeness, and contiguity of these assemblies. To generate *de novo* assemblies, we use minimap2 [108] to report all read overlaps and miniasm [48] to construct the assembly from these overlaps. We use miniasm because it allows us to observe the effect of the reads on the assemblies without performing additional error correction steps on input reads [113] and their final assembly [35]. To measure the assembly accuracy, we use dnadiff [114] to evaluate 1) the portion of the reference genome that can align to a given assembly (i.e., Genome Fraction), 2) the average identity of assemblies (i.e., Average Identity) when compared to their respective reference genomes, and 3) insertions and deletions of nucleotides (or bases) in the sequence when compared to a reference or other sequences. (i.e., Total Indels and Indel Ratio (%)). Total Indels represents the sum of all the insertions and deletions in the assembled sequence when compared to a reference or other sequences. The Indel Ratio is a measure of the relative abundance of indels compared to the total length of the assembled sequence (calculated using Total Indels / Assembly Length) × 100. This metric helps to understand the proportion of the assembly that contains insertions and deletions. To measure statistics related to the contiguity and completeness of the assemblies, such as the overall assembly length, average GC content (i.e., the ratio of G and C bases in an assembly), and NG50 statistics (i.e., shortest contig at the half of the overall reference genome length), we use QUAST [115]. We assume that the reference genomes are high-quality representative of the sequenced samples that we basecall the reads from when comparing assemblies to their corresponding reference genomes. The higher the values of the average identity, genome fraction, and NG50 results, the higher the quality of the assembly and, hence the better the corresponding basecaller. When the values of the average GC and assembly length results are closer to that of the corresponding reference genome, the better the assembly and the corresponding basecaller. We use Inspector [116] to calculate the overall quality value (QV) of an assembly. The QV score is determined by considering structural and small-scale errors in proportion to the total number of base pairs in the assemblies. High-quality sequences have higher QV scores, indicating a low probability of sequencing errors, while low-quality sequences have lower QV scores, suggesting a higher likelihood of errors.

### Read mapping

We basecall the raw electrical signals into reads using each of the subject basecallers. We map the resulting read set to the reference genome of the same species using the state-of-the-art read mapper, minimap2 [108]. We use the default parameter values for mapping ONT reads using the preset parameter *-x map-ont*. We use the *stats* tool from the SAMtools library [117] to obtain four key statistics on the quality of read mapping results, the total number of mismatches, the total number of mapped bases, the total number of mapped reads, and the total number of unmapped reads. We normalize the total number of base mismatches and the total number of mapped bases using the total number of bases in the reads, while for the total number of mapped reads and the total number of unmapped reads, we normalize using the total number of reads.

## 6. Data Availability

The read set used in this study was downloaded from https://bridges.monash.edu/articles/dataset/Raw_fast5s/7676174, while the reference set can be downloaded from https://bridges.monash.edu/articles/dataset/Reference_genomes/7676135. For the human genome, we download reads from https://labs.epi2me.io/gm24385_2020.11/, while the reference genome is available at https://github.com/marbl/HG002. All trained models and generated reads can be downloaded from https://zenodo.org/record/7413316. We ensure unbiased, fair, and consistent evaluation by retraining all the basecallers using the official ONT dataset [63].

## 7. Code Availability

Source code with the instructions for reproducing the results is publicly available at: https://github.com/Xilinx/neuralArchitectureReshaping. Scripts used to perform basecalling accuracy analysis are available at: https://github.com/rrwick/Basecalling-comparison.

## 8. Acknowledgments

We thank the SAFARI Research Group members for their valuable feedback and the stimulating intellectual and scholarly environment they provide. SAFARI Research Group acknowledges the generous gifts of their industrial partners, including Google, Huawei, Intel, Microsoft, VMware, and Xilinx. This research was partially supported by the Semiconductor Research Corporation. SAFARI Research Group acknowledges support from the European Union’s Horizon programme for research and innovation under grant agreement No. 101047160, project BioPIM. Special thanks to Alessandro Pappalardo for his support with quantization-aware training. We appreciate valuable discussions with Giovanni Mariani. Thanks to AMD for providing access to the HPC fund cluster [118].

## 9. Author Contributions

G.S., M.A., K.D., and A.K. conceived `RUBICON`. G.S. designed and implemented `RUBICON`. G.S., C.F., and M.C. collected data and performed the evaluations. K.D., H.C., and O.M. supervised the work. G.S., M.A., K.D., and C.F. wrote the manuscript. All authors reviewed and edited the manuscript. All authors analyzed the results. All authors read and approved the final manuscript.

## 10. Competing Interests

All authors declare no competing interests.

## Supplementary Material for

### S1. Quantization-Aware Basecaller Architecture Search (QABAS)

`QABAS` automates the process of finding efficient and high-performance hardware-aware genomics base-callers. Figure S1 shows the workflow overview of `QABAS`. The raw sequencing data 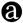 is provided as input to `QABAS`, which can be obtained through sequencing a new sample, downloading from publicly-available databases, or computer simulation. `QABAS` uses such a set of data as training (𝔻_*train*_) and evaluation set (𝔻_*eval*_) while automatically designing a basecaller. To achieve a basecaller design that provides high throughput, we add hardware constraints 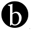, in terms of latency or throughput, to `QABAS`. A hardware-aware basecaller can better use the underlying hardware features and greatly accelerate inference speed. As a result, it improves the overall basecalling efficiency.

**Figure S1:**
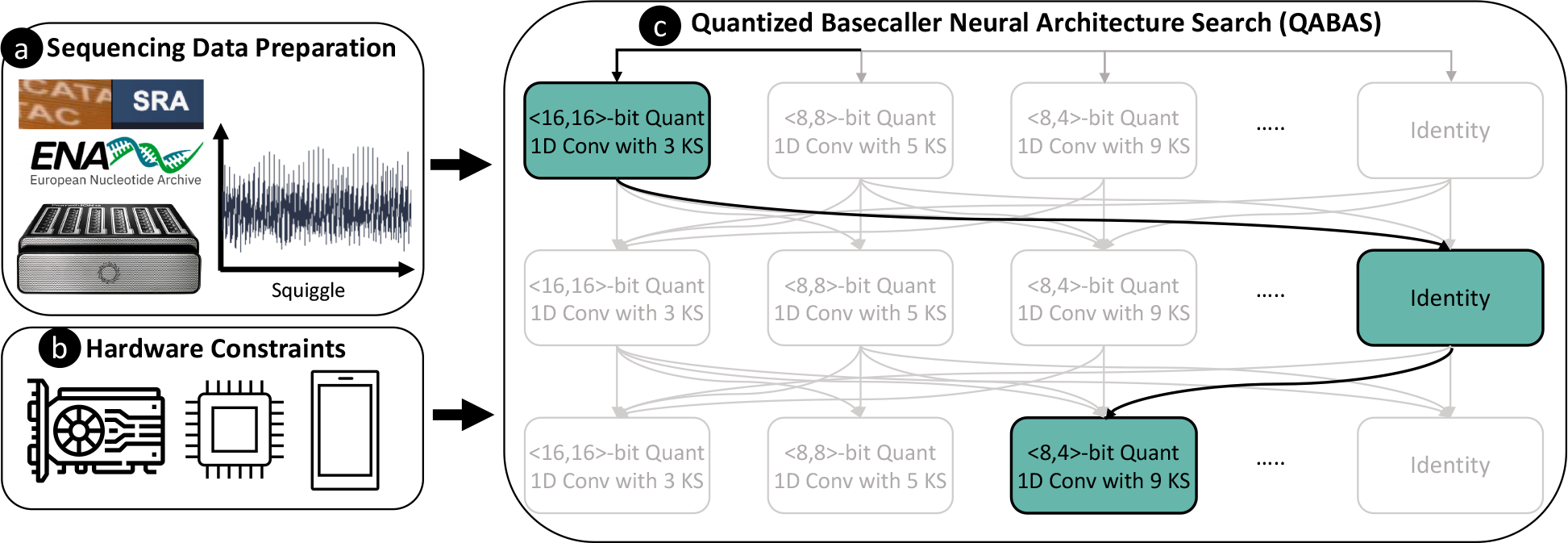
Overview of QABAS. QABAS evaluates a different set of candidate operations for convolution (conv) and quantization bits. In the figure, we show different options for kernel size (KS) (e.g., 3, 5, 9, etc.) and quantization bits (4-b, 8-b, and 16-b) for each network layer. The identity operator removes a layer to get a shallower network.

`QABAS` 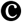 leverages automated machine learning (AutoML) algorithms [57] using neural architecture search (NAS) to design an efficient hardware basecaller by exploring and evaluating different neural network architectures from a pre-defined search space. The search space consists of the possible neural ℳnetwork architectural options while 𝕄 ∈ ℳ is a sub-architecture from 𝕄. The goal is to find an optimal sub-architecture 𝕄^∗^ using Equation S1 that minimizes the training loss (ℒ _*train*_) while going over 𝔻_*train*_ and gives maximum accuracy with the 𝔻_*eval*_.

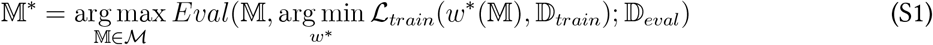

where *w*^∗^(𝕄) represent the weights of sub-architecture 𝕄^∗^.

#### QABAS Search Space

We define the search spaceℳ as sufficiently large to enable a powerful neural architecture search. A larger space enables the search algorithm to cover more architectures to increase the chance of finding a powerful architecture. However, a larger search space makes converging more difficult for the search algorithm.

Our model search space has sequentially connected blocks, where each block receives input from its direct previous block. We formulate the NAS problem for hardware-aware genomics basecaller as finding: (a) the computational operations in each basic block^3^ of a basecaller, including operations in a skip connection block, and (b) quantization bit-width for weights and activations for each neural network layer to perform low-precision computation. Quantization is the reduction of the bit-width precision at which calculations are performed in a neural network to reduce memory and computational complexity. Adding quantization exploration dramatically increases the model search space (∼6.72× 10^20^ additional viable options in our search space). However, performing a joint search for computational blocks and quantization bits is crucial because: (1) optimizing these two components in separate stages could lead to sub-optimal results as the best network architecture for the full-precision model is not necessarily the optimal one after quantization, and (2) independent exploration would also require considerable search time and energy consumption because of many viable design options [121]. Therefore, `QABAS` searches for both the computational operations present in each basic block of a basecaller and the quantization bits used by these computational operations. In doing so, we tailor the neural network architecture and computation to align with the hardware’s capabilities.

#### QABAS Search Algorithm

`QABAS` evaluates different neural network architectures using differentiable neural architecture search (DNAS) [122–124]. DNAS follows a weight-sharing approach of reusing weights of previously optimized architectures from the neural architecture search space. For example, if sub-architecture 𝕄_1_ has only one additional layer compared to sub-architecture 𝕄_2_. In such a scenario, 𝕄_1_ can use most weights from 𝕄_2_. Therefore, the search procedure gets accelerated in DNAS compared to training each sub-architecture individually.

DNAS formulates the entire search space as a super-network and distills a target network from this super-network. Traditional NAS approaches [57] often sample many different architectures from the search space and train each architecture from scratch to validate its performance. Such an approach requires heavy computational resources that could lead to thousands of GPU hours of overhead. One way to overcome this issue is to use NAS with heuristic-based methods [125, 126], such as genetic algorithms that select individual architectures from the current *population* to be *parents* and uses them to produce the *children* for the next generation. However, such methods still suffer from the problem of retraining each sample architecture from scratch. Therefore, DNAS provides an efficient solution by sharing computation among different architectures, as many of them have similar properties.

In `QABAS`, we construct an over-parameterized super-network with all possible candidate options. The super-network shares weights among sub-architecture. During the search phase, `QABAS` searches for the optimal: (a) architectural parameter *α*: likelihood that a computational operation will be preserved in the final architecture; and (b) network weights *w*: weights of convolution layers. We use ProxylessNAS [127] to binarize the architectural parameter (i.e., *α*∈ {0,1}) to reduce memory consumption during the search phase. At the end of the search phase, the operators with the highest architectural weight are preserved, while others are eliminated. Since the NAS search procedure is focused on optimizing the super-network, the final sub-network architecture 𝕄∗, with all the preserved operations, is retrained to convergence to fully optimize its network weights.

#### Quantization-Aware Hardware Metric

Current state-of-the-art basecallers [29, 30, 40–43, 45, 73] are hardware-agnostic. They only focus on improving the accuracy without paying attention to its inference efficiency. For example, Fast-bonito [42] uses NAS for basecalling architecture search, however, it does not consider any hardware-related metrics during the architecture search. Therefore, such approaches lead to over-provisioned basecallers with a large number of parameters and model sizes that are unoptimized for mixed-precision computation (see Section 2.1). We overcome this inefficiency in `QABAS` by adding hardware constraints, in terms of inference latency, to the `QABAS` search phase. Thus, `QABAS` aims to find an efficient neural network architecture for basecalling that is also optimized for hardware implementation. During the search process, `QABAS` sequentially selects a sub-network from the super-network. The expected latency of the sub-network is the sum of the latencies of each operation in the network. Before the start of the `QABAS` search phase, we profile the latencies of operations present in the search space on targeted hardware to build a latency estimator. We also incorporate the latency while using different quantization bit-widths for the weights and activations in our latency estimator. This latency estimator is utilized to guide the `QABAS` search process.

`QABAS`’s objective function (ℒ_QABAS_) minimizes a joint cross-entropy error to: (a) provide better basecalling accuracy by minimizing the training loss (ℒ_*train*_) while going over 𝔻_*train*_, and (b) minimize a regularization term (ℒ_*reg*_) to find a sub-network 𝕄 with inference latency (𝕃_𝕄_) that satisfies our inference latency constraints. We add latency constraints by using a target latency parameter (𝕃_*tar*_) to the regularization term ℒ_*reg*_ to guide the search process. For example, in case we want a small model, then we can provide a higher 𝕃_*tar*_ value, or vice versa.

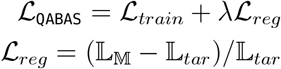

where *λ* is a parameter to control the tradeoff between the basecalling accuracy and the model latency. As different hardware provides different latencies for the same layers chosen from the `QABAS` search space, the user can customize the `RUBICON` framework for their target hardware by adjusting hardware-specific parameters (i.e., using the `applied_hardware` flag in `RUBICON` [128]). We provide an additional `reference_latency` flag in `QABAS` to guide the search of basecalling architecture to find an architecture that meets certain latency constraints. This coupling of target hardware latency ensures the basecaller architecture is finely tuned to operate optimally on the intended hardware. We provide an example latency estimator for our target hardware (i.e., AIE) in `RUBICON`. However, our integration with the open-source nn-Meter [99] tool allows users to freely configure hardware settings through the `applied_hardware` flag in `RUBICON`. This integration enhances adaptability, enabling efficient optimization and deployment across different hardware environments.

### S2. SkipClip: Skip Connection Removal by Teaching

Deep neural networks often rely on skip connections to address vanishing gradient problems during training [53]. Skip connections provide a direct path for error propagation, allowing gradients to flow without vanishing [129]. Additionally, they prevent saturation issues in deep neural networks, making them more effective. Similarly, deep learning-based basecallers [29, 30, 40–43, 45, 73] use skip connections to mitigate the vanishing gradient and saturation problems. However, adding skip connections introduces the following three issues for hardware acceleration. First, skip connections increases the data-lifetime. The layers whose activations are reused in subsequent layers must wait for this activation reuse (or buffer the activations in memory) before accepting new input and continuing to compute. This leads to high resource and storage requirements due to data duplication. Second, they introduce irregularity in neural network architecture as these connections span across non-adjacent layers. Third, skip connections require additional computation to adjust the channel size to match the channel size at the non-consecutive layer’s input. Thus, increasing model parameters and model size. Therefore, networks without skip connections have more regular topologies that translate better to hardware acceleration.

To address these issues, we propose `SkipClip`, a first skip connection remover for basecallers. `SkipClip` gradually removes skip connections using knowledge distillation (KD) [58, 64], where a pretrained larger model (teacher) guides a smaller model (student) to maintain performance without skip connections. As shown in Figure S2, `SkipClip` starts with a pretrained over-parameterized model as the teacher, which is not updated during the training of the student network. We use our final `QABAS` model as the student network. We achieve skip removal by letting the teacher teach the student to perform well on basecalling. At the start of every training epoch, `SkipClip` removes a skip connection from a block, starting from the input side, while performing KD. This is done until all skip connections are removed from the student network. `SkipClip` gets the best of both worlds: a highly accurate and topologically regular neural network without skip connections.

During the `SkipClip`, we perform a forward pass of both the student and the teacher model, while we perform a backward pass only for the student model to update its weights. The loss to update the student network’s weight during the backward pass ℒ_SkipClip_) is calculated with Equation S2, where we use a weighing of the actual student loss (ℒ_*S*_) and distillation loss (ℒ_*D*_) using an alpha (*α*) hyper-parameter. The student and the teacher model compute probabilities *f*_*T*_ and *f*_*S*_ for output labels (i.e., nucleotides A, C, G, T) in the forward pass, respectively. We use cross entropy (ℒ_*CR*_) in the probability distributions to calculate the distillation loss (ℒ_*D*_) as in Equation S3. The temperature (*τ*) variable is used for *softening* the probability distributions, i.e., it controls the weight of knowledge from the teacher network for a student network to absorb. As we raise the *τ*, the resulting soft label probability distribution becomes richer in information.

**Figure S2:**
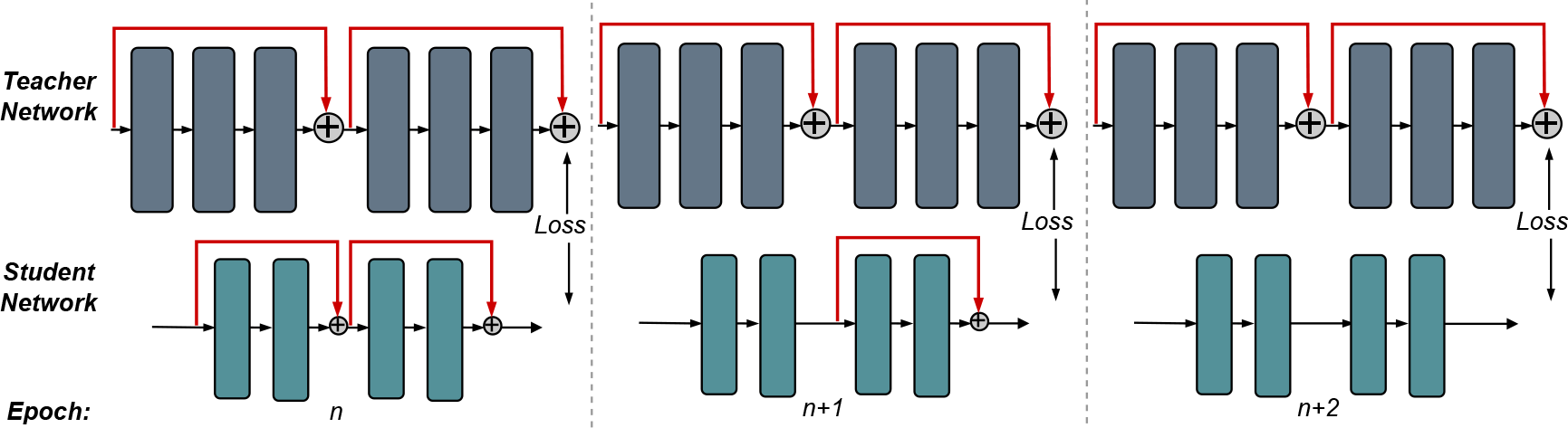
Overview of SkipClip process for three epochs. We start with a large, overprovisioned floating-point precision model as the teacher network and our QABAS mixed-precision model as the student network. During the training, SkipClip removes a skip connection from the student network every n epoch, starting with the first skip connection encountered in the network from the input.

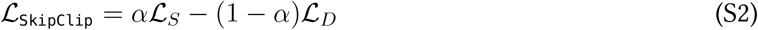

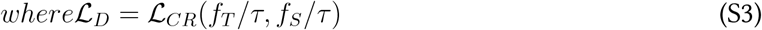

#### S2.1. Sensitivity to Skip Connection

Many state-of-the-art deep learning-based basecallers [29, 30, 40–43, 45, 73] incorporate skip connections to improve their basecalling accuracy. Figure S3 shows the accuracy of `Bonito_CTC` using two different configurations of skip connections (`s1` and `s2`) and one configuration without any skip connections (`s3`) and compares it to the baseline `Bonito_CTC` architecture. In `s1` configuration, we reduce the number of repeats in each block to one, while in `s2` configuration, we use only one block with maximum channel size, maximum kernel size, and the maximum number of repeats.

**Figure S3:**
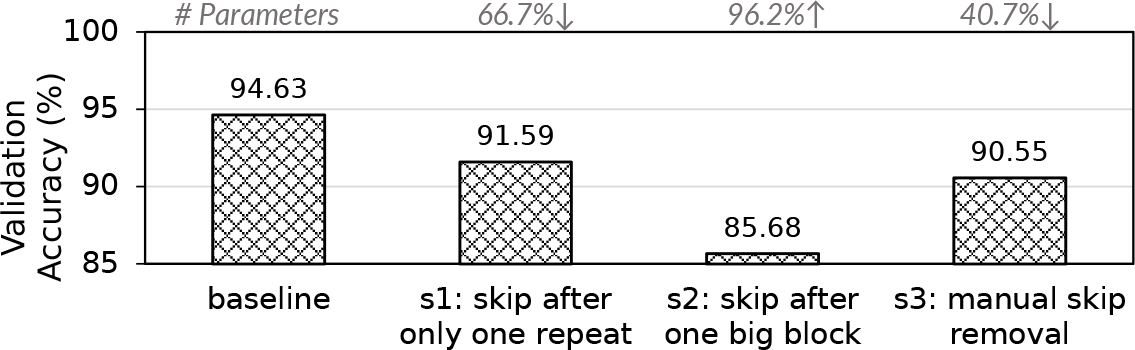
Basecaller sensitivity to skip connections.

For `s3` configuration, we manually remove all the skip connections from each block in `Bonito_CTC`. We also annotate the change in model parameters compared to the baseline model. `Bonito_CTC` architecture comprises several blocks, each consisting of a time channel separable convolution sub-block (referred to as repeat). We make two major observations. First, the number of sub-blocks we provide skip connection plays an important role. In `s1` configuration, we observe that by using only one repeat, we reduce the accuracy by 2.84% with 66.7% lower model parameters, while by merging all the blocks into one big block in `s2` configuration, we observe 8.75% lower accuracy with 96.2% higher model parameters. Second, manually removing all the skip connections in `s3` configuration leads to 40.7% lower model parameters at the expense of a 3.88% loss in accuracy. This performance degradation is because, during neural network training, these connections provide a direct path for propagating the error through the layers and dealing with the vanishing gradient problem, allowing deep networks to learn properly and converge during training. Therefore, manual removal of skip connections can lead to lower basecalling performance. We conclude that skip connections are critical for basecalling accuracy.

#### S2.2. Hyper-parameter Tuning for SkipClip

In Figure S4, we show the effect of two critical hyper-parameters of `SkipClip` (alpha (*α*) and temperature (*τ*)) on the validation accuracy of `Bonito_CTC`. We observe that as we raise *α* while keeping *τ* constant, the basecaller accuracy increases. At higher *α*, `SkipClip` gives more importance to the student loss than the distillation loss during the backward pass. We use *α* =0.9 throughout our experiments.

**Figure S4:**
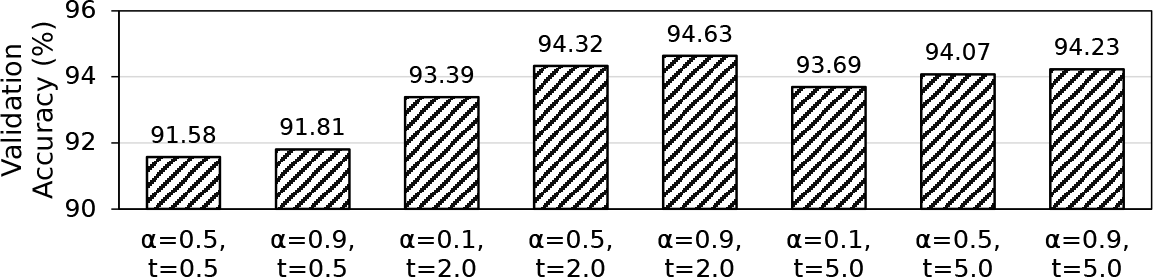
Sensitivity of SkipClip to hyper-parameters alpha (*α*) and temperature (*τ*).

For *τ*, we experiment with values ranging from 0.5 to 5.0. Increasing *τ* provides more knowledge from the teacher network for a student network to absorb. We observe at *τ* =2, `SkipClip` provides the highest accuracy. Further increasing *τ* does not provide benefits because the student network cannot absorb knowledge provided by the teacher network.

### S3. RUBICALL Architecture

Figure S5 shows the architecture of `RUBICALL`. We develop `RUBICALL` using `QABAS` and `SkipClip`. The `RUBICALL` architecture is composed of 28 quantized convolution blocks containing ∼3.3 million model parameters. Each block consists of quantized grouped 1-dimensional convolution and quantized pointwise 1-dimensional convolution where every layer is quantized to a different domain. The convolution operation is followed by batch normalization (Batch Norm) [119] and a quantized rectified linear unit (QuantReLU) [120] activation function. The final output is passed through a connectionist temporal classification (CTC) [130] layer to produce the decoded sequence of nucleotides (A, C, G, T). CTC is used to provide the correct alignment between the input and the output sequence.

In a learning task, 𝒳represents feature space with label 𝒴, where a machine learning model is responsible for estimating a function *f* : 𝒳 → 𝒴. RUBICALL first splits a long read in electrical-signal format (e.g., millions of signals) into multiple smaller chunks (e.g., thousands of samples per chunk) and then basecalls these chunks. `RUBICALL` uses the input signal (or squiggle) as 𝒳to predict nucleotides as label 𝒴. The CTC layer assigns a probability for all possible labels in 𝒴given an at 𝒳each time-step. The nucleotide with the highest probability is selected as the final output.

**Figure S5:**
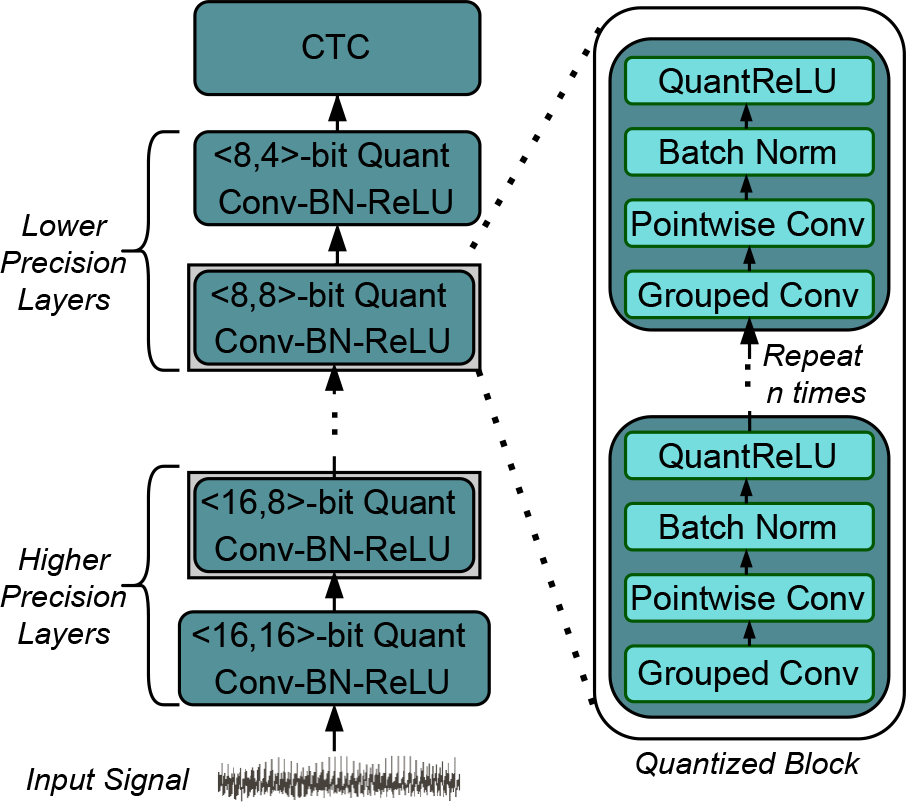
Overview of RUBICALL architecture. The normalized input signal is passed through a succession of quantized convolution blocks. Each block is composed of several processing steps (convolution, batch normalization, and activation). We represent the quantization as a tuple *<*weight, activation*>*. Initial layers use a higher precision for weights and activations, while the final layers use a lower precision. The final output is passed through a connectionist temporal classification (CTC) to produce the decoded sequence of nucleotides.

### S4. Comparison to More Accurate Basecallers

Our goal is to make basecalling highly efficient and fast by building the first framework for specializing and optimizing machine learning-based basecaller. Currently, we focus on CNN-based basecallers because: (1) they are the most widely used basecallers, and (2) the fundamental multiply-accumulate (MAC) operation in a CNN model is amenable to hardware acceleration, unlike the operations in RNN-based basecallers. As `Bonito_CTC` has the same backend as `RUBICALL` (i.e., Quartznet [50]), we consider it as an expert-designed model. `Bonito_CRF`’s super high accuracy (`Bonito_CRF-sup`) model is an RNN-based basecaller that provides more accuracy than `Bonito_CRF-fast` at the expense of a much larger model. We compare the overall basecalling throughput of `RUBICALL` with that of the baseline basecallers in terms of basecalling accuracy, model parameters, and model size in Figure S6(a), S6(b), and S6(c), respectively.

**Figure S6:**
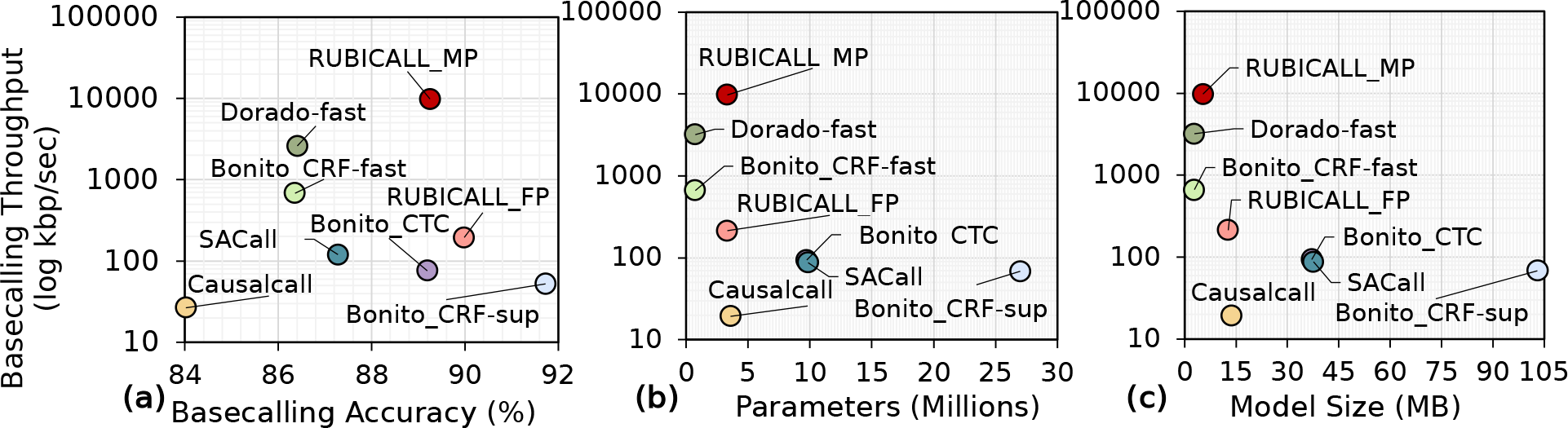
Comparison of average basecalling throughput for RUBICALL-MP with baseline basecaller in terms of: (a) average basecalling accuracy, (b) model parameters, and (c) model size.

**Table S1:**
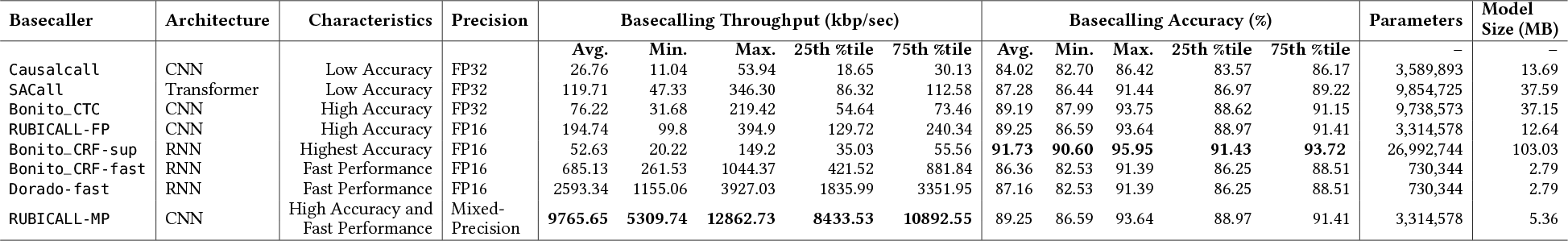
Comparison of RUBICALL-MP with baseline basecallers in terms of model architecture, characteristics, precision, basecalling throughput, basecalling accuracy, parameters, and model size. For basecalling throughput and basecalling accuracy, we report average (Avg.), minimum (Min.), maximum (Max.), 25th percentile (25th %tile), and 75th percentile (75th %tile) values for all the basecallers.

In addition to our previous observations from Figure 5, we make three new observations from Figure S6 and Table S1. First, `RUBICALL-MP` has 185.54× the performance of the highly-accurate `Bonito_CRF-sup`. `RUBICALL-MP` is the only basecaller that provides both higher performance and accuracy when compared to all the other evaluated basecallers. Second, `Bonito_CRF-sup` uses 7.52×, 36.96×, 2.77×, 2.74×, 36.96 ×, and 8.14 model parameters leading to a model size of 7.53×, 36.93×, 2.77×, and 19.22× compared to `Causalcall`, `Bonito_CRF-fast`, `Bonito_CTC`, `SACall`, `Dorado-fast` and `RUBICALL-MP`, respectively. Third, `Bonito_CRF-sup` is 5.37% more accurate than its throughout-optimized version, `Bonito_CRF-fast`, which provides up to 13.02× higher basecalling performance. We conclude that the high accuracy of a basecaller comes at a substantial cost in terms of lower throughput due to the higher number of model parameters and model size.

### S5. Evaluation on Other Hardware Platforms

We also evaluate the performance of `RUBICALL` and all the other basecallers on NVIDIA A40 [70] GPU with 48GiB DRAM and AMD EPYC 7442 [87] 24-Core with 256GiB DRAM. Compared to the AMD MI210 [69], the NVIDIA A40 has a 1.65× higher peak compute performance while maintaining a 2.35× lower peak memory bandwidth.

We make two major observations from Figure S7. First, `RUBICALL-MP` on AIE consistently outper-forms A40 by 502.52×, 14.67×, 104.14×, 111.25×, 45.61×, and 3.19× higher performance compared to `Causalcall`, `Bonito_CRF-fast`, `Bonito_CTC`, `SACall`, `RUBICALL-FP`, and `Dorado-fast`, respectively. Second, for compute-bound basecallers, A40 provides 1.23×, 1.18×, and 1.09× higher performance than AMD MI210 (Figure 6) for `Bonito_CTC`, `Dorado-fast`, and `RUBICALL-FP`, respectively. For memory-bound basecallers, A40 provides 1.38×, 1.03×, and 1.36× lower performance for `Causalcall`, `Bonito_CRF-fast`, and `SACall`, respectively. We conclude that `RUBICON` provides benefits across multiple hardware platforms.

**Figure S7:**
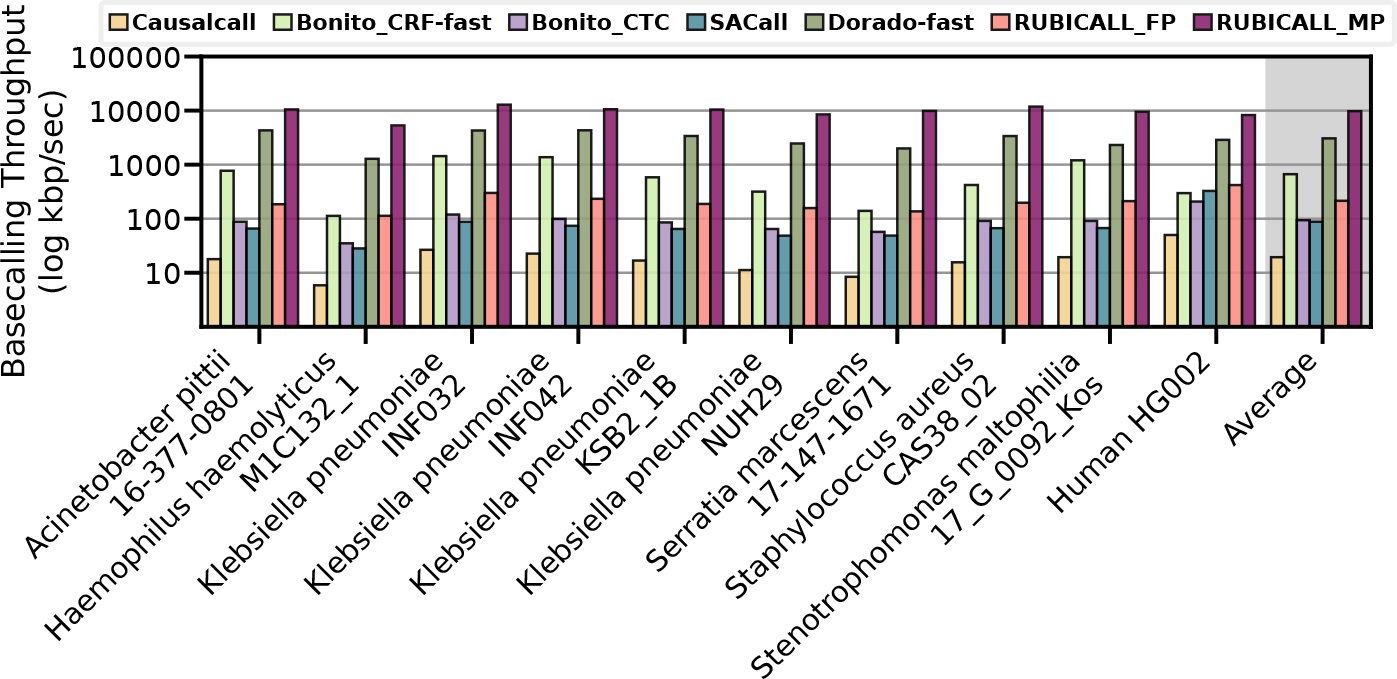
Performance comparison of RUBICALL (using floating-point precision (RUBICALL-FP) and mixed-precision (RUBICALL-MP)) and five state-of-the-art basecallers on NVIDIA A40 [70]. The y-axis is on a logarithmic scale.

### S6. Analysis of Mapped Reads and Mapped Bases

Table S2 shows the average read length, the overall number of mapped reads, the number of mapped bases, and the ratio of mapped bases to the mapped reads. Our goal is to evaluate the tools in terms of the read lengths they can generate and the alignable fraction of these reads to their corresponding reference genomes. We make three key observations. First, we find that the average read lengths are similar across different basecallers for each dataset, except `Causalcall` for the human genome. This indicates that the substantial differences in read length are unlikely to influence the ratio of mapped bases to the number of mapped reads, while the number of alignable sequences within each read and the number of mapped reads can have the main effect on such a ratio. Second, we find that basecallers provide a similar number of mapped reads and the ratio of mapped bases to the mapped reads for each dataset, except `Causalcall` for the human genome. These similarities mainly indicate that the unalignable reads and the unalignable regions within each read are likely to be similar across basecallers, leading to similar ratios of mapped bases to mapped reads when mapping reads with similar average read lengths. Third, we find that `Causalcall` provides exceptions for the human genome in terms of the average read length and the mapped bases to the mapped reads ratio. This is mainly because `Causalcall` fails to basecall all raw signals for the human genome and provides a subset of basecalled reads that other basecallers generate, leading to inaccurate analysis overall. We conclude that almost all basecallers, except `Causalcall`, generate reads with similar average read lengths and reads with similar alignable regions, although these similarities differ by certain percentages’ as we discuss in Section 2.5.2.

**Table S2:** Read mapping comparison of RUBICALL with baseline basecallers in terms of mean length of individual sequencing reads in a dataset (Avg. Length), the total number of mapped reads (Mapped Reads), the total number of mapped bases (Mapped Bases), and the ratio of total number of mapped reads to mapped bases.

| Dataset | Basecaller | Avg. Length | Mapped Reads | Mapped Bases | #Mapped Bases/<br>#Mapped Reads |
| --- | --- | --- | --- | --- | --- |
| Acinetobacter<br>pittii 16-377-0801 | causalcall | 25,718.6 | 4,434 | 114,159,528 | 25,746.4 |
|  | Bonito_CRF-fast | 26,151.3 | 4,452 | 110,907,740 | 24,911.9 |
|  | Bonito_CTC | 24,879.1 | 4,457 | 110,183,466 | 24,721.4 |
|  | SACall | 25,153.3 | 4,451 | 111,997,940 | 25,162.4 |
|  | Dorado-fast | 26,151.1 | 4,452 | 110,907,740 | 24,911.9 |
|  | RUBICALL | 25,000.8 | 4,452 | 111,405,897 | 25,023.8 |
| Haemophilus<br>haemolyticus<br>M1C132_1 | causalcall | NA | NA | NA | NA |
|  | Bonito_CRF-fast | 9,835.7 | 6,444 | 64,816,196 | 10,058.4 |
|  | Bonito_CTC | 8,862.8 | 6,201 | 73,573,092 | 11,864.7 |
|  | SACall | 6,871.1 | 4,028 | 43,233,160 | 10,733.2 |
|  | Dorado-fast | 9,844.8 | 6,444 | 64,816,196 | 10,058.4 |
|  | RUBICALL | 7,751.5 | 6,287 | 63,415,299 | 10,086.7 |
| Klebsiella<br>pneumoniae<br>INF032 | causalcall | 35,781.9 | 15,150 | 542,123,428 | 35,783.7 |
|  | Bonito_CRF-fast | 36,556.4 | 15,147 | 533,045,454 | 35,191.5 |
|  | Bonito_CTC | 35,189.1 | 15,152 | 519,659,064 | 34,296.4 |
|  | SACall | 35,078.5 | 15,150 | 531,456,488 | 35,079.6 |
|  | Dorado-fast | 36,624.7 | 15,147 | 533,045,454 | 35,191.5 |
|  | RUBICALL | 35,420.7 | 15,153 | 536,724,002 | 35,420.3 |
| Klebsiella<br>pneumoniae<br>INF042 | causalcall | 48,483.8 | 11,236 | 542,123,428 | 48,248.8 |
|  | Bonito_CRF-fast | 49,617.5 | 11,252 | 533,045,454 | 47,373.4 |
|  | Bonito_CTC | 46,198.4 | 11,273 | 519,659,064 | 46,097.7 |
|  | SACall | 46,298.3 | 11,198 | 531,456,488 | 47,459.9 |
|  | Dorado-fast | 49,621.4 | 11,252 | 533,045,454 | 47,373.4 |
|  | RUBICALL | 46,637.6 | 11,268 | 536,724,002 | 47,632.6 |
| Klebsiella<br>pneumoniae<br>KSB2_1B | causalcall | 24,039.6 | 16,642 | 401,041,491 | 24,098.2 |
|  | Bonito_CRF-fast | 24,723.9 | 16,744 | 384,436,100 | 22,959.6 |
|  | Bonito_CTC | 22,918.8 | 16,803 | 385,157,295 | 22,921.9 |
|  | SACall | 22,917.5 | 16,371 | 381,266,978 | 23,289.2 |
|  | Dorado-fast | 24,728.0 | 16,744 | 384,436,100 | 22,959.6 |
|  | RUBICALL | 23,141.9 | 16,783 | 388,897,351 | 23,172.1 |
| Klebsiella<br>pneumoniae<br>NUH29 | causalcall | 16,233.7 | 14,954 | 243,112,795 | 16,257.4 |
|  | Bonito_CRF-fast | 16,435.7 | 15,056 | 229,123,038 | 15,218.1 |
|  | Bonito_CTC | 15,182.0 | 15,152 | 233,135,041 | 15,386.4 |
|  | SACall | 15,536.5 | 15,088 | 234,764,649 | 15,559.7 |
|  | Dorado-fast | 16,419.0 | 15,056 | 229,123,038 | 15,218.1 |
|  | RUBICALL | 15,300.8 | 15,113 | 231,523,267 | 15,319.5 |
| Serratia<br>marcescens<br>17-147-1671 | causalcall | 8,198.0 | 12,729 | 104,864,058 | 8,238.2 |
|  | Bonito_CRF-fast | 8,456.2 | 16,667 | 133,916,776 | 8,034.8 |
|  | Bonito_CTC | 8,024.8 | 16,715 | 133,754,055 | 8,002.0 |
|  | SACall | 8,167.1 | 16,665 | 136,289,479 | 8,178.2 |
|  | Dorado-fast | 8,465.1 | 16,667 | 133,916,776 | 8,034.8 |
|  | RUBICALL | 8,076.5 | 16,696 | 134,916,360 | 8,080.8 |
| Staphylococcus<br>aureus<br>CAS38_02 | causalcall | 21,425.2 | 11,038 | 236,529,129 | 21,428.6 |
|  | Bonito_CRF-fast | 21,932.6 | 11,047 | 237,020,597 | 21,455.7 |
|  | Bonito_CTC | 21,455.7 | 11,047 | 232,091,657 | 21,009.5 |
|  | SACall | 21,372.6 | 11,047 | 236,103,648 | 21,372.6 |
|  | Dorado-fast | 21,930.8 | 11,047 | 237,020,597 | 21,455.7 |
|  | RUBICALL | 21,501.0 | 11,047 | 237,521,476 | 21,501.0 |
| Stenotrophomonas<br>maltophilia<br>17_G_0092_Kos | causalcall | 31,415.3 | 15,946 | 501,018,868 | 31,419.7 |
|  | Bonito_CRF-fast | 31,736.6 | 15,959 | 470,408,299 | 29,476.1 |
|  | Bonito_CTC | 29,453.8 | 15,997 | 474,913,094 | 29,687.6 |
|  | SACall | 30,095.7 | 15,985 | 481,168,698 | 30,101.3 |
|  | Dorado-fast | 31,727.6 | 15,959 | 470,408,299 | 29,476.1 |
|  | RUBICALL | 29,676.7 | 15,980 | 474,401,853 | 29,687.2 |
| Human<br>HG002 | causalcall | 11,201.7 | 163,984 | 2,612,902,733 | 15,933.9 |
|  | Bonito_CRF-fast | 37,755.9 | 238,205 | 10,627,000,000 | 44,612.8 |
|  | Bonito_CTC | 35,831.5 | 243,686 | 10,212,000,000 | 41,906.4 |
|  | SACall | 32,815.7 | 203,228 | 9,080,197,989 | 44,679.9 |
|  | Dorado-fast | 37,991.9 | 238,474 | 10,627,000,000 | 44,562.5 |
|  | RUBICALL | 37,141.2 | 245,373 | 10,591,000,000 | 43,162.9 |

### S7. K-mer Counting Analysis

We analyze the occurrence of k-mer (i.e., substrings of length k) in a given sequence of basecalled reads and their assemblies in Figure S8 and Figure S9, respectively. We use BBMap [131] to collect the number of unique k-mers and the frequency of each unique k-mer in a given sequence. During our analysis, we vary the value of k from 15 to 31. Based on our empirical analysis, we set the k value for our evaluated bacterial species to 15, where we observe distinct peaks of unique k-mers. We do not perform k-mer frequency analysis for the human genome due to the low coverage of the human genome in our experiments. We make the following two observations from Figure S8 and Figure S9. First, `RUBICALL` has distinct peaks for all the evaluated species, often matching the k-mer composition generated from `Bonito_CTC`. Second, `Bonito_CRF-fast` and `Dorado-fast` generate similar k-mer compositions as they both have the same neural network architecture.

**Figure S8:**
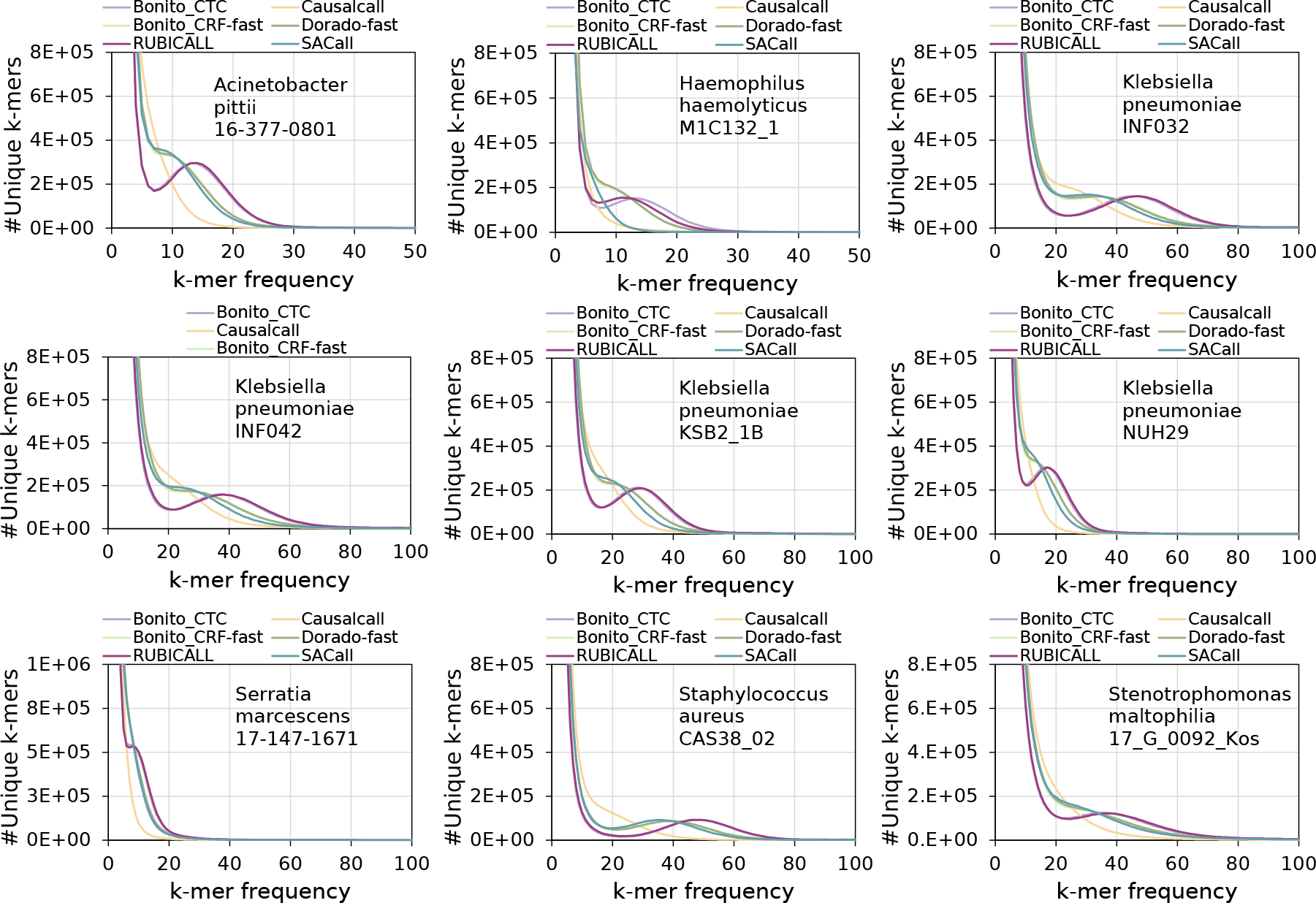
K-mer frequency analysis of generated reads from RUBICALL and all the other evaluated basecallers.

**Figure S9:**
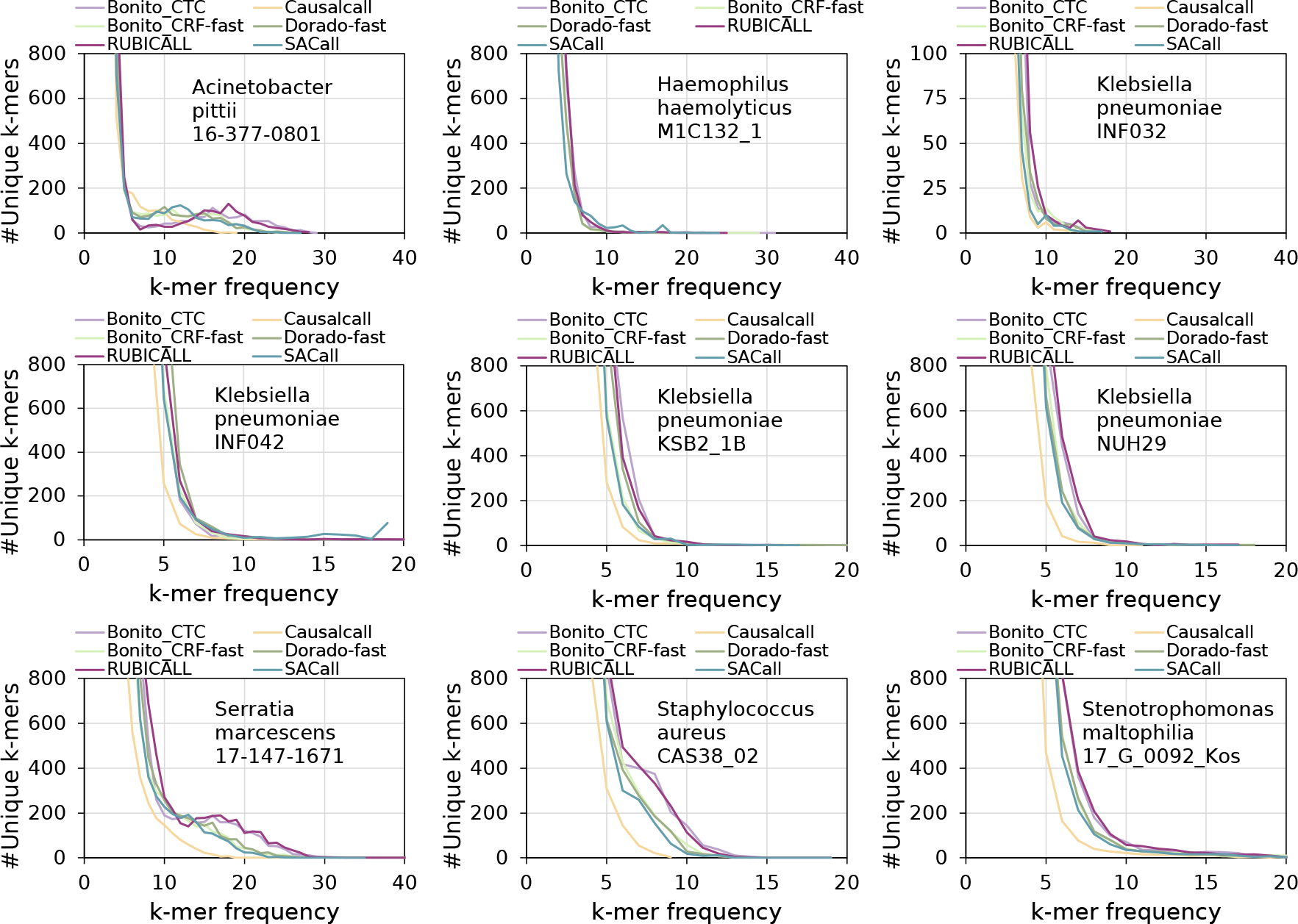
K-mer frequency analysis of generated assemblies of reads from RUBICALL and all the other evaluated basecallers.

Table S3 presents an analysis of k-mer frequencies in the raw reads and the corresponding assemblies. We include common sequences and read-to-assembly ratios to provide a comprehensive view of the similarities and disparities in sequence representation, aiding in assessing data quality and the performance of the assembly algorithms. We observe that the k-mers identified as over-represented in the assemblies are mainly observed as over-represented k-mers in read sets for most basecallers. These over-represented k-mers are likely to appear due to the particular repetitive regions of each genome, making k-mers appear a larger amount of times for these regions. Therefore, there is potentially no additional insertion or depletion of these k-mers during the assembly process.

**Table S3:** Comparison of under and over-represented sequences (k-mers) in reads and assemblies for all the evaluated basecallers. For both under and over-represented sequences, we show common sequences (Common) and the ratio of k-mer frequencies between reads and assemblies (Ratio).

| Dataset | Basecaller | Under-Represented |  |  |  | Over-Represented |  |  |  |
| --- | --- | --- | --- | --- | --- | --- | --- | --- | --- |
|  |  | Read | Assembly | Common | Ratio | Read | Assembly | Common | Ratio |
| Acinetobacter pitti<br>16-377-0801 | Causalcall | 73,096,681 | 3,768,504 | 3,221,668 | 0.855 | 6,263 | 187 | 187 | 1.000 |
|  | Bonito.CRF-fast | 55,359,526 | 3,747,817 | 2,367,806 | 0.632 | 17,983 | 360 | 360 | 1.000 |
|  | Bonito.CTC | 44,790,782 | 3,593,419 | 1,814,762 | 0.505 | 29,097 | 296 | 296 | 1.000 |
|  | SACall | 55,660,535 | 3,625,236 | 2,381,232 | 0.657 | 15,534 | 368 | 368 | 1.000 |
|  | Dorado-fast | 55,775,603 | 3,760,029 | 2,404,108 | 0.639 | 18,137 | 425 | 425 | 1.000 |
|  | RUBICALL | 44,085,891 | 3,609,296 | 1,793,772 | 0.497 | 30,316 | 430 | 430 | 1.000 |
| Haemophilus haemolyticus<br>M1C132_1 | Causalcall | 31,021,572 | NA | NA | NA | 1,552 | NA | NA | NA |
|  | Bonito.CRF-fast | 42,355,232 | 2,077,823 | 2,076,203 | 0.999 | 2,865 | 54 | 53 | 0.981 |
|  | Bonito.CTC | 35,847,257 | 1,919,667 | 700,997 | 0.365 | 33,713 | 94 | 94 | 1.000 |
|  | SACall | 36,998,888 | 2,000,357 | 1,998,679 | 0.999 | 22,517 | 98 | 61 | 0.622 |
|  | Dorado-fast | 41,939,332 | 2,074,317 | 2,072,740 | 0.999 | 4,714 | 37 | 37 | 1.000 |
|  | RUBICALL | 33,917,316 | 1,929,302 | 1,127,668 | 0.584 | 23,221 | 100 | 100 | 1.000 |
| Klebsiella pneumoniae<br>INF032 | Causalcall | 197,327,230 | 4,850,481 | 3,833,326 | 0.790 | 15,891 | 6 | 6 | 1.000 |
|  | Bonito.CRF-fast | 169,124,267 | 4,914,464 | 3,353,593 | 0.682 | 35,175 | 22 | 22 | 1.000 |
|  | Bonito.CTC | 155,835,445 | 4,758,242 | 2,718,894 | 0.571 | 53,149 | 20 | 20 | 1.000 |
|  | SACall | 176,768,581 | 4,747,776 | 3,366,011 | 0.709 | 28,038 | 12 | 12 | 1.000 |
|  | Dorado-fast | 167,899,392 | 4,916,810 | 3,351,520 | 0.682 | 35,059 | 24 | 24 | 1.000 |
|  | RUBICALL | 150,348,189 | 4,776,357 | 2,689,193 | 0.563 | 62,356 | 26 | 26 | 1.000 |
| Klebsiella pneumoniae<br>INF042 | Causalcall | 211,565,073 | 5,171,081 | 4,222,032 | 0.816 | 22,204 | 3 | 3 | 1.000 |
|  | Bonito.CRF-fast | 178,074,237 | 5,212,872 | 3,532,743 | 0.678 | 61,454 | 29 | 29 | 1.000 |
|  | Bonito.CTC | 162,568,221 | 4,964,190 | 2,843,401 | 0.573 | 81,948 | 36 | 36 | 1.000 |
|  | SACall | 186,186,165 | 5,024,228 | 3,609,853 | 0.718 | 40,440 | 41 | 41 | 1.000 |
|  | Dorado-fast | 174,755,139 | 5,508,965 | 3,808,628 | 0.691 | 63,644 | 23 | 23 | 1.000 |
|  | RUBICALL | 158,433,298 | 4,989,328 | 2,818,523 | 0.565 | 92,045 | 37 | 37 | 1.000 |
| Klebsiella pneumoniae<br>KSB2_1B | Causalcall | 180,267,220 | 5,064,568 | 4,648,903 | 0.918 | 8,844 | 6 | 6 | 1.000 |
|  | Bonito.CRF-fast | 152,878,553 | 5,122,808 | 4,044,261 | 0.789 | 23,755 | 16 | 16 | 1.000 |
|  | Bonito.CTC | 137,461,268 | 4,859,560 | 3,174,990 | 0.653 | 27,211 | 14 | 14 | 1.000 |
|  | SACall | 158,736,471 | 4,903,831 | 4,117,531 | 0.840 | 16,090 | 12 | 12 | 1.000 |
|  | Dorado-fast | 150,458,414 | 5,265,881 | 4,164,287 | 0.791 | 24,938 | 19 | 19 | 1.000 |
|  | RUBICALL | 134,066,854 | 4,876,789 | 3,186,025 | 0.653 | 31,451 | 17 | 17 | 1.000 |
| Klebsiella pneumoniae<br>NUH29 | Causalcall | 140,405,375 | 5,060,601 | 5,004,375 | 0.989 | 835 | 1 | 1 | 1.000 |
|  | Bonito.CRF-fast | 110,315,181 | 5,060,829 | 4,696,878 | 0.928 | 4,320 | 22 | 22 | 1.000 |
|  | Bonito.CTC | 97,405,757 | 4,775,503 | 4,182,794 | 0.876 | 5,177 | 20 | 20 | 1.000 |
|  | SACall | 112,996,097 | 4,844,709 | 4,613,301 | 0.952 | 2,941 | 16 | 16 | 1.000 |
|  | Dorado-fast | 108,585,877 | 5,043,253 | 4,679,036 | 0.928 | 4,483 | 23 | 23 | 1.000 |
|  | RUBICALL | 95,201,166 | 4,789,453 | 4,136,482 | 0.864 | 5,645 | 25 | 25 | 1.000 |
| Serratia marcescens<br>17-147-1671 | Causalcall | 66,514,376 | 5,334,807 | 5,193,034 | 0.973 | 30,238 | 4 | 4 | 1.000 |
|  | Bonito.CRF-fast | 63,963,265 | 5,399,858 | 4,554,600 | 0.843 | 61,321 | 4 | 4 | 1.000 |
|  | Bonito.CTC | 53,413,056 | 5,217,101 | 3,821,412 | 0.732 | 61,275 | 9 | 9 | 1.000 |
|  | SACall | 64,337,585 | 5,284,226 | 4,534,343 | 0.858 | 54,872 | 1 | 1 | 1.000 |
|  | Dorado-fast | 63,535,166 | 5,451,215 | 4,583,027 | 0.841 | 62,006 | 4 | 4 | 1.000 |
|  | RUBICALL | 51,724,568 | 5,243,385 | 3,741,711 | 0.714 | 64,943 | 9 | 9 | 1.000 |
| Staphylococcus aureus<br>CAS38_02 | Causalcall | 106,477,765 | 2,791,690 | 2,784,563 | 0.997 | 3,375 | 3 | 3 | 1.000 |
|  | Bonito.CRF-fast | 72,908,007 | 2,813,307 | 2,774,962 | 0.986 | 12,170 | 33 | 33 | 1.000 |
|  | Bonito.CTC | 59,047,673 | 2,736,120 | 2,669,644 | 0.976 | 18,475 | 37 | 37 | 1.000 |
|  | SACall | 73,372,106 | 2,741,321 | 2,710,663 | 0.989 | 10,487 | 29 | 29 | 1.000 |
|  | Dorado-fast | 73,708,021 | 2,824,623 | 2,786,854 | 0.987 | 11,902 | 37 | 37 | 1.000 |
|  | RUBICALL | 58,939,315 | 2,739,823 | 2,675,209 | 0.976 | 18,186 | 84 | 84 | 1.000 |
| Stenotrophomonas maltophilia<br>17_G_0092_Kos | Causalcall | 183,625,102 | 4,653,477 | 4,000,323 | 0.860 | 25,103 | 132 | 132 | 1.000 |
|  | Bonito.CRF-fast | 144,980,026 | 4,647,752 | 3,083,896 | 0.664 | 127,258 | 210 | 210 | 1.000 |
|  | Bonito.CTC | 127,334,549 | 4,383,078 | 2,495,090 | 0.569 | 170,398 | 285 | 285 | 1.000 |
|  | SACall | 139,640,543 | 4,422,404 | 3,045,239 | 0.689 | 112,124 | 201 | 201 | 1.000 |
|  | Dorado-fast | 143,087,662 | 4,594,794 | 3,087,855 | 0.672 | 127,653 | 201 | 201 | 1.000 |
|  | RUBICALL | 121,924,168 | 4,391,308 | 2,430,608 | 0.554 | 197,883 | 327 | 327 | 1.000 |

## Footnotes

1 A skip connection allows to skip some of the layers in the neural network and feeds the output of one layer as the input to the next layers.

2 We define knee point as the point beyond which a basecaller is unable to basecall at an acceptable level of accuracy.

3 Our basic block consists of one-dimensional (1-D) convolution, batch normalization [119], and rectified linear unit (ReLU) [120].

## Notes

https://bridges.monash.edu/articles/dataset/Raw_fast5s/7676174

https://bridges.monash.edu/articles/dataset/Reference_genomes/7676135

https://labs.epi2me.io/gm24385_2020.11/

https://github.com/marbl/HG002

### Competing Interest Statement

The authors have declared no competing interest.

### Summary of Updates

Added new experiments. Updated writing and streamlined text.

## References

[1] Ginsburg G, Phillips K. Precision Medicine: From Science To Value. Health Affairs. 2018 05;37:694–701.

[2] Aryan Z, Szanto A, Pantazi A, Reddi T, Rheinstein C, Powers W, et al. Moving Genomics to Routine Care: An Initial Pilot in Acute Cardiovascular Disease. Circulation Genomic and precision medicine. 2020 October;13(5):406–416. Available from: 10.1161/CIRCGEN.120.002961.

[3] Clark MM, Hildreth A, Batalov S, Ding Y, Chowdhury S, Watkins K, et al. Diagnosis of Genetic Diseases in Seriously Ill Children by Rapid Whole-Genome Sequencing and Automated Phenotyping and Interpretation. Science translational medicine. 2019 April;11(489):eaat6177. Available from: 10.1126/scitranslmed.aat6177.

[4] Kingsmore SF, Smith LD, Kunard CM, Bainbridge M, Batalov S, Benson W, et al. A Genome Sequencing System for Universal Newborn Screening, Diagnosis, and Precision Medicine for Severe Genetic Diseases. American journal of human genetics. 2022 September;109(9):1605—1619. Available from: 10.1016/j.ajhg.2022.08.003.

[5] Ginsburg GS, Willard HF. Genomic and Personalized Medicine: Foundations and Applications. Translational Research. 2009;154(6):277–87. Special Issue on Personalized Medicine. Available from: https://www.sciencedirect.com/science/article/pii/S1931524409002746.

[6] Bloom JS, Sathe L, Munugala C, Jones EM, Gasperini M, Lubock NB, et al. Massively Scaled-Up Testing for SARS-CoV-2 RNA via Next-Generation Sequencing of Pooled and Barcoded Nasal and Saliva Samples. Nature Biomedical Engineering. 2021 Jul;5(7):657–65. Available from: 10.1038/s41551-021-00754-5.

[7] Quick J, Loman NJ, Duraffour S, Simpson JT, Severi E, Cowley L, et al. Real-Time, Portable Genome Sequencing for Ebola Surveillance. Nature Research; 2016-02-11.

[8] Yelagandula R, Bykov A, Vogt A, Heinen R, Özkan E, Strobl MM, et al. Multiplexed Detection of SARS-CoV-2 and Other Respiratory Infections in High Throughput by SARSeq. Nature communications. 2021 May;12(1):3132. Available from: https://europepmc.org/articles/PMC8149640.

[9] Le VTM, Diep BA. Selected Insights from Application of Whole-Genome Sequencing for Outbreak Investigations. Current Opinion in Critical Care. 2013;19:432–439.

[10] Nikolayevskyy V, Kranzer K, Niemann S, Drobniewski F. Whole Genome Sequencing of M.tuberculosis for Detection of Recent Transmission and Tracing Outbreaks: A Systematic Review. Tuberculosis. 2016 03;98.

[11] Meyer F, Fritz A, Deng ZL, Koslicki D, Gurevich A, Robertson G, et al. Critical Assessment of Metagenome Interpretation-The Second Round of Challenges. bioRxiv. 2021.

[12] LaPierre N, Alser M, Eskin E, Koslicki D, Mangul S. Metalign: Efficient Alignment-Based Metagenomic Profiling via Containment Min Hash. Genome biology. 2020;21(1):1–15.

[13] LaPierre N, Mangul S, Alser M, Mandric I, Wu N, Koslicki D, et al. MiCoP: Microbial Community Profiling Method for Detecting Viral and Fungal Organisms in Metagenomic Samples. BMC Genomics. 2019 06;20:423.

[14] Meyer F, Fritz A, Deng ZL, Koslicki D, Lesker TR, Gurevich A, et al. Critical Assessment of Metagenome Interpretation: The Second Round of Challenges. Nature Methods. 2022 Apr;19(4):429–40. Available from: 10.1038/s41592-022-01431-4.

[15] Pollard MO, Gurdasani D, Mentzer AJ, Porter T, Sandhu MS. Long Reads: Their Purpose and Place. Human Molecular Genetics. 2018 Aug;27(R2):R234–41.

[16] Senol Cali D, Kim JS, Ghose S, Alkan C, Mutlu O. Nanopore Sequencing Technology and Tools for Genome Assembly: Computational Analysis of the Current State, Bottlenecks and Future Directions. Briefings in Bioinformatics. 2019 Jul;20(4):1542–59.

[17] Amarasinghe SL, Su S, Dong X, Zappia L, Ritchie ME, Gouil Q. Opportunities and Challenges in Long-Read Sequencing Data Analysis. Genome biology. 2020;21(1):1–16.

[18] Logsdon GA, Vollger MR, Eichler EE. Long-Read Human Genome Sequencing and its Applications. Nature Reviews Genetics. 2020;21(10):597–614.

[19] Wang Y, Zhao Y, Bollas A, Wang Y, Au KF. Nanopore Sequencing Technology, Bioinformatics and Applications. Nature biotechnology. 2021;39(11):1348–65.

[20] Jain M, Koren S, Miga KH, Quick J, Rand AC, Sasani TA, et al. Nanopore Sequencing and Assembly of a Human Genome with Ultra-Long Reads. Nature Biotechnology. 2018 Apr;36(4):338–45. Available from: 10.1038/nbt.4060.

[21] Gong L, Wong CH, Idol J, Ngan CY, Wei CL. Ultra-Long Read Sequencing for Whole Genomic DNA Analysis. JoVE. 2019 Mar;(145):e58954. Available from: https://www.jove.com/t/58954.

[22] Branton D, Deamer DW, Marziali A, Bayley H, Benner SA, Butler T, et al. The potential and challenges of nanopore sequencing. Nature biotechnology. 2008;26(10):1146–53.

[23] Van Dijk EL, Jaszczyszyn Y, Naquin D, Thermes C. The Third Revolution in Sequencing Technology. Trends in Genetics. 2018;34(9):666–81.

[24] Ardui S, Ameur A, Vermeesch JR, Hestand MS. Single Molecule Real-Time (SMRT) Sequencing Comes of Age: Applications and Utilities for Medical Diagnostics. Nucleic acids research. 2018;46(5):2159–68.

[25] Jain M, Koren S, Miga KH, Quick J, Rand AC, Sasani TA, et al. Nanopore Sequencing and Assembly of a Human Genome with Ultra-Long Reads. Nature Biotechnology. 2018 Apr;36(4):338–45.

[26] Kchouk M, Gibrat JF, Elloumi M. Generations of Sequencing Technologies: From First to Next Generation. Biology and Medicine. 2017;9(3).

[27] Weirather JL, de Cesare M, Wang Y, Piazza P, Sebastiano V, Wang XJ, et al. Comprehensive Comparison of Pacific Biosciences and Oxford Nanopore Technologies and Their Applications to Transcriptome Analysis. F1000Research. 2017;6.

[28] Jain M, Olsen HE, Paten B, Akeson M. The Oxford Nanopore MinION: Delivery of Nanopore Sequencing to the Genomics Community. Genome biology. 2016;17(1):1–11.

[29] Wick RR, Judd LM, Holt KE. Performance of Neural Network Basecalling Tools for Oxford Nanopore Sequencing. Genome biology. 2019;20(1):1–10.

[30] Pages-Gallego M, de Ridder J. Comprehensive and Standardized Benchmarking of Deep Learning Architectures for Basecalling Nanopore Sequencing Data. bioRxiv. 2022.

[31] Alser M, Lindegger J, Firtina C, Almadhoun N, Mao H, Singh G, et al. From Molecules to Genomic Variations: Accelerating Genome Analysis via Intelligent Algorithms and Architectures. Computational and Structural Biotechnology Journal. 2022.

[32] Alser M, Rotman J, Deshpande D, Taraszka K, Shi H, Baykal PI, et al. Technology Dictates Algorithms: Recent Developments in Read Alignment. Genome Biology. 2021 Aug;22(1):249.

[33] Zhang Yz, Akdemir A, Tremmel G, Imoto S, Miyano S, Shibuya T, et al. Nanopore Basecalling from a Perspective of Instance Segmentation. BMC bioinformatics. 2020.

[34] Dias R, Torkamani A. Artificial Intelligence in Clinical and Genomic Diagnostics. Genome medicine. 2019;11(1):1–12.

[35] Firtina C, Kim JS, Alser M, Senol Cali D, Cicek AE, Alkan C, et al. Apollo: A Sequencing-Technology-Independent, Scalable and Accurate Assembly Polishing Algorithm. Bioinformatics. 2020 Jun;36(12):3669–79.

[36] Rang FJ, Kloosterman WP, de Ridder J. From Squiggle to Basepair: Computational Approaches for Improving Nanopore Sequencing Read Accuracy. Genome Biology. 2018 Jul;19(1):90. Available from: 10.1186/s13059-018-1462-9.

[37] Mao H, Alser M, Sadrosadati M, Firtina C, Baranwal A, Cali DS, et al. Genpip: In-memory acceleration of genome analysis via tight integration of basecalling and read mapping. In: 2022 55th IEEE/ACM International Symposium on Microarchitecture (MICRO). IEEE; 2022. p. 710–26.

[38] Lv X, Chen Z, Lu Y, Yang Y. An End-to-End Oxford Nanopore Basecaller Using Convolution-Augmented Transformer. In: 2020 IEEE International Conference on Bioinformatics and Biomedicine (BIBM). IEEE; 2020. p. 337–42.

[39] Zeng J, Cai H, Peng H, Wang H, Zhang Y, Akutsu T. Causalcall: Nanopore Basecalling Using a Temporal Convolutional Network. Frontiers in Genetics. 2020:1332.

[40] Perešíni P, BoŽa V, Brejová B, Vinař T. Nanopore Base Calling on the Edge. Bioinformatics. 2021;37(24):4661–7.

[41] Lou Q, Janga SC, Jiang L. Helix: Algorithm/Architecture Co-design for Accelerating Nanopore Genome Base-calling. In: Proceedings of the ACM International Conference on Parallel Architectures and Compilation Techniques; 2020. p. 293–304.

[42] Xu Z, Mai Y, Liu D, He W, Lin X, Xu C, et al. Fast-bonito: A Faster Deep Learning Based Basecaller for Nanopore Sequencing. Artificial Intelligence in the Life Sciences. 2021;1:100011.

[43] Konishi H, Yamaguchi R, Yamaguchi K, Furukawa Y, Imoto S. Halcyon: An Accurate Basecaller Exploiting an Encoder– Decoder Model with Monotonic Attention. Bioinformatics. 2021;37(9):1211–7.

[44] Huang N, Nie F, Ni P, Luo F, Wang J. SACall: A Neural Network Basecaller for Oxford Nanopore Sequencing Data Based on Self-Attention Mechanism. IEEE/ACM Transactions on Computational Biology and Bioinformatics. 2020.

[45] Neumann D, Reddy AS, Ben-Hur A. RODAN: A Fully Convolutional Architecture for Basecalling Nanopore RNA Sequencing Data. BMC bioinformatics. 2022;23(1):1–9.

[46] NVIDIA. NVIDIA A10 Tensor Core GPU, https://www.nvidia.com/en-us/data-center/products/a10-gpu/;.

[47] Benchmarking the Oxford Nanopore Technologies basecallers on AWS, https://aws.amazon.com/blogs/hpc/benchmarking-the-oxford-nanopore-technologies-basecallers-on-aws/;.

[48] Li H. Minimap and Miniasm: Fast Mapping and De Novo Assembly for Noisy Long Sequences. Bioinformatics. 2016 Jul;32(14):2103–10.

[49] Ulrich JU, Lutfi A, Rutzen K, Renard BY. ReadBouncer: precise and scalable adaptive sampling for nanopore sequencing. Bioinformatics. 2022;38.

[50] Kriman S, Beliaev S, Ginsburg B, Huang J, Kuchaiev O, Lavrukhin V, et al. QuartzNet: Deep Automatic Speech Recognition with 1D Time-Channel Separable Convolutions. In: ICASSP 2020-2020 IEEE International Conference on Acoustics, Speech and Signal Processing (ICASSP). IEEE; 2020. p. 6124–8.

[51] Majumdar S, Balam J, Hrinchuk O, Lavrukhin V, Noroozi V, Ginsburg B. Citrinet: Closing the Gap Between Non-Autoregressive and Autoregressive End-to-End Models for Automatic Speech Recognition. arXiv preprint arXiv:210401721. 2021.

[52] Gulati A, Qin J, Chiu CC, Parmar N, Zhang Y, Yu J, et al. Conformer: Convolution-Augmented Transformer for Speech Recognition. arXiv preprint arXiv:200508100. 2020.

[53] Szegedy C, Ioffe S, Vanhoucke V, Alemi AA. Inception-v4, Inception-ResNet and the Impact of Residual Connections on Learning. In: Thirty-first AAAI conference on artificial intelligence; 2017. .

[54] Singh G, Diamantopoulos D, Stuijk S, Hagleitner C, Corporaal H. Low precision processing for high order stencil computations. In: International Conference on Embedded Computer Systems. Springer; 2019. p. 403–15.

[55] Singh G, Diamantopoulos D, Hagleitner C, Gómez-Luna J, Stuijk S, Mutlu O, et al. NERO: A near high-bandwidth memory stencil accelerator for weather prediction modeling. In: 2020 30th International Conference on Field-Programmable Logic and Applications (FPL). IEEE; 2020. p. 9–17.

[56] Singh G. Designing, Modeling, and Optimizing Data-Intensive Computing Systems. arXiv preprint arXiv:220808886. 2022.

[57] Zoph B, Le QV. Neural Architecture Search with Reinforcement Learning. arXiv preprint arXiv:161101578. 2016.

[58] Buciluă C, Caruana R, Niculescu-Mizil A. Model Compression. In: Proceedings of the 12th ACM SIGKDD international conference on Knowledge discovery and data mining; 2006. p. 535–41.

[59] LeCun Y, Denker J, Solla S. Optimal Brain Damage. Advances in neural information processing systems. 1989;2.

[60] Han S, Mao H, Dally WJ. Deep Compression: Compressing Deep Neural Networks with Pruning, Trained Quantization and Huffman Coding. arXiv. 2015.

[61] Han S, Pool J, Tran J, Dally W. Learning both Weights and Connections for Efficient Neural Network. Advances in neural information processing systems. 2015;28.

[62] Frankle J, Carbin M. The Lottery Ticket Hypothesis: Finding Sparse, Trainable Neural Networks. arXiv preprint arXiv:180303635. 2018.

[63] Bonito, https://github.com/nanoporetech/bonito;.

[64] Hinton G, Vinyals O, Dean J, et al. Distilling the Knowledge in a Neural Network. arXiv preprint arXiv:150302531. 2015;2(7).

[65] Versal ACAP AI Core Series Product Selection Guide, https://www.xilinx.com/content/dam/xilinx/support/documents/selection-guides/versal-ai-core-product-selection-guide.pdf;.

[66] Kruschke JK, Movellan JR. Benefits of Gain: Speeded Learning and Minimal Hidden Layers in Back-Propagation Networks. IEEE Transactions on systems, Man, and Cybernetics. 1991;21(1):273–80.

[67] Liu Z, Sun M, Zhou T, Huang G, Darrell T. Rethinking the Value of Network Pruning. arXiv preprint arXiv:181005270. 2018.

[68] Gale T, Elsen E, Hooker S. The State of Sparsity in Deep Neural Networks. arXiv preprint arXiv:190209574. 2019.

[69] AMD. AMD Instinct MI210 Accelerator, https://www.amd.com/system/files/documents/amd-instinct-mi210-brochure.pdf;.

[70] NVIDIA. NVIDIA A40, https://images.nvidia.com/content/Solutions/data-center/a40/nvidia-a40-datasheet.pdf;.

[71] Ferrarini M, Moretto M, Ward JA, Šurbanovski N, Stevanović V, Giongo L, et al. An Evaluation of the PacBio RS Platform for Sequencing and De Novo Assembly of a Chloroplast Genome. BMC Genomics. 2013 Oct;14(1):670.

[72] Chen YC, Liu T, Yu CH, Chiang TY, Hwang CC. Effects of GC Bias in Next-Generation-Sequencing Data on De Novo Genome Assembly. PLOS ONE. 2013 Apr;8(4):e62856.

[73] Zhang Z, Park CY, Theesfeld CL, Troyanskaya OG. An Automated Framework for Efficiently Designing Deep Convolutional Neural Networks in Genomics. Nature Machine Intelligence. 2021;3(5):392–400.

[74] Singh G, Gómez-Luna J, Mariani G, Oliveira GF, Corda S, Stuijk S, et al. Napel: Near-memory computing application performance prediction via ensemble learning. In: 2019 56th ACM/IEEE Design Automation Conference (DAC). IEEE; 2019. p. 1–6.

[75] Singh G, Nadig R, Park J, Bera R, Hajinazar N, Novo D, et al. Sibyl: Adaptive and Extensible Data Placement in Hybrid Storage Systems Using Reinforcement Learning. In: Proceedings of the 49th Annual International Symposium on Computer Architecture. ISCA ‘22. New York, NY, USA: Association for Computing Machinery; 2022. p. 320–336. Available from: 10.1145/3470496.3527442.

[76] Nurvitadhi E, Sim J, Sheffield D, Mishra A, Krishnan S, Marr D. Accelerating Recurrent Neural Networks in Analytics Servers: Comparison of FPGA, CPU, GPU, and ASIC. In: FPL; 2016. .

[77] Singh G, Alser M, Senol Cali D, Diamantopoulos D, Gómez-Luna J, Corporaal H, et al. FPGA-Based Near-Memory Acceleration of Modern Data-Intensive Applications. IEEE Micro. 2021 Aug;41(4):39–48.

[78] Singh G, Khodamoradi A, Denolf K, Lo J, Gomez-Luna J, Melber J, et al. SPARTA: Spatial Acceleration for Efficient and Scalable Horizontal Diffusion Weather Stencil Computation. In: Proceedings of the 37th International Conference on Supercomputing; 2023. p. 463–76.

[79] Singh G, Diamantopoulos D, Gómez-Luna J, Hagleitner C, Stuijk S, Corporaal H, et al. Accelerating weather prediction using near-memory reconfigurable fabric. ACM Transactions on Reconfigurable Technology and Systems (TRETS). 2022;15(4):1–27.

[80] Umuroglu Y, Fraser NJ, Gambardella G, Blott M, Leong P, Jahre M, et al. Finn: A framework for fast, scalable binarized neural network inference. In: Proceedings of the 2017 ACM/SIGDA international symposium on field-programmable gate arrays; 2017. .

[81] Alipanahi B, Delong A, Weirauch MT, Frey BJ. Predicting the Sequence Specificities of DNA-and RNA-Binding Proteins by Deep Learning. Nature biotechnology. 2015;33(8):831–8.

[82] Boemo MA. DNAscent v2: Detecting Replication Forks in Nanopore Sequencing Data with Deep Learning. BMC genomics. 2021;22(1):1–8.

[83] Sabba S, Smara M, Benhacine M, Hameurlaine A. Residual Neural Network for Predicting Super-Enhancers on Genome Scale. In: International Conference on Artificial Intelligence and its Applications. Springer; 2021. p. 32–42.

[84] Barnes GH, Brown RM, Kato M, Kuck DJ, Slotnick DL, Stokes RA. The ILLIAC IV Computer. IEEE Transactions on Computers. 1968.

[85] Open Neural Network Exchange (ONNX), https://github.com/onnx/onnx;.

[86] Baskin C, Liss N, Schwartz E, Zheltonozhskii E, Giryes R, Bronstein AM, et al. Uniq: Uniform noise injection for non-uniform quantization of neural networks. ACM Transactions on Computer Systems (TOCS). 2021;37(1-4):1–15.

[87] Introducing 3rd Gen AMD EPYC™ Processors, https://www.amd.com/en/events/epyc;.

[88] Tullsen DM, Eggers SJ, Levy HM. Simultaneous Multithreading: Maximizing On-Chip Parallelism. In: ISCA; 1995. .

[89] RDIMM, https://www.micron.com/products/dram-modules/rdimm;.

[90] Ubuntu 20.04.3 LTS (Focal Fossa), https://releases.ubuntu.com/20.04/;.

[91] GCC, the GNU Compiler Collection;. Available from: https://gcc.gnu.org/.

[92] AMD. ROCm, https://github.com/RadeonOpenCompute/ROCm;.

[93] NVIDIA System Management Interface, https://developer.nvidia.com/nvidia-system-management-interface;.

[94] NVIDIA CUDA Compiler Driver NVCC, https://docs.nvidia.com/cuda/cuda-compiler-driver-nvcc/index.html;.

[95] ARM Cortex-A72 MPCore Processor Technical Reference Manual r0p3, https://developer.arm.com/documentation/100095/0003;.

[96] Kraken 2, https://github.com/DerrickWood/kraken2;.

[97] Larsen ACM, Knudsen CA, Hansen MN. Palamut - An Expansion of the Bonito basecaller using language models [Master’s thesis]; 2020. Available at https://projekter.aau.dk/projekter/files/334904330/MI104F20_Speciale Paper__21_.pdf.

[98] NNI, https://github.com/microsoft/nni;.

[99] nn Meter Team MR. nn-Meter: Towards Accurate Latency Prediction of Deep-Learning Model Inference on Diverse Edge Devices; 2021. Available from: https://github.com/microsoft/nn-Meter.

[100] Pappalardo A. Xilinx/brevitas. Zenodo; 2021. Available from: 10.5281/zenodo.3333552.

[101] Kingma DP, Ba J. Adam: A Method for Stochastic Optimization. arXiv. 2014.

[102] KLDivLoss, https://pytorch.org/docs/stable/generated/torch.nn.KLDivLoss.html;.

[103] PyTorch, https://pytorch.org/;.

[104] TORCH.NN, https://pytorch.org/docs/stable/nn.html;.

[105] ONT. Dorado, https://github.com/nanoporetech/dorado.git;.

[106] PyTorch C++ API, https://pytorch.org/cppdocs/;.

[107] Silvestre-Ryan J, Holmes I. Pair consensus decoding improves accuracy of neural network basecallers for nanopore sequencing. Genome biology. 2021;22:1–6.

[108] Li H. Minimap2: Pairwise Alignment for Nucleotide Sequences. Bioinformatics. 2018 Sep;34(18):3094–100.

[109] Rebaler, https://github.com/rrwick/Rebaler;.

[110] Vaser R, Sović I, Nagarajan N, Šikić M. Fast and accurate de novo genome assembly from long uncorrected reads. Genome Research. 2017.

[111] Robertson G, Schein J, Chiu R, Corbett R, Field M, Jackman SD, et al. De Novo Assembly and Analysis of RNA-seq Data. Nature methods. 2010;7(11):909–12.

[112] Li B, Ruotti V, Stewart RM, Thomson JA, Dewey CN. RNA-Seq Gene Expression Estimation with Read Mapping Uncertainty. Bioinformatics. 2010;26(4):493–500.

[113] Firtina C, Bar-Joseph Z, Alkan C, Cicek AE. Hercules: A Profile HMM-based Hybrid Error Correction Algorithm for Long Reads. Nucleic Acids Research. 2018 Nov;46(21):e125–5.

[114] Marçais G, Delcher AL, Phillippy AM, Coston R, Salzberg SL, Zimin A. MUMmer4: A Fast and Versatile Genome Alignment System. PLOS Computational Biology. 2018 Jan;14(1):e1005944.

[115] Gurevich A, Saveliev V, Vyahhi N, Tesler G. QUAST: Quality Assessment Tool for Genome Assemblies. Bioinformatics. 2013 Apr;29(8):1072–5.

[116] Chen Y, Zhang Y, Wang AY, Gao M, Chong Z. Accurate long-read de novo assembly evaluation with Inspector. Genome Biology. 2021;22(1):1–21.

[117] Li H, Handsaker B, Wysoker A, Fennell T, Ruan J, Homer N, et al. The Sequence Alignment/Map (SAM) Format and SAMtools. Bioinformatics. 2009;25(16):2078–9.

[118] AMD HPC Fund, https://www.amd.com/en/corporate/hpc-fund;.

[119] Ioffe S, Szegedy C. Batch Normalization: Accelerating Deep Network Training by Reducing Internal Covariate Shift. In: International conference on machine learning. PMLR; 2015. p. 448–56.

[120] Agarap AF. Deep Learning Using Rectified Linear Units (ReLU). arXiv. 2018.

[121] Ren P, Xiao Y, Chang X, Huang PY, Li Z, Chen X, et al. A Comprehensive Survey of Neural Architecture Search: Challenges and Solutions. ACM Computing Surveys (CSUR). 2021;54(4):1–34.

[122] Liu H, Simonyan K, Yang Y. Darts: Differentiable Architecture Search. arXiv. 2018.

[123] Luo R, Tian F, Qin T, Chen E, Liu TY. Neural Architecture Optimization. Advances in neural information processing systems. 2018;31.

[124] Xie L, Chen X, Bi K, Wei L, Xu Y, Wang L, et al. Weight-Sharing Neural Architecture Search: A Battle to Shrink the Optimization Gap. ACM Computing Surveys (CSUR). 2021;54(9):1–37.

[125] Real E, Moore S, Selle A, Saxena S, Suematsu YL, Tan J, et al. Large-Scale Evolution of Image Classifiers. In: International Conference on Machine Learning. PMLR; 2017. p. 2902–11.

[126] Xie L, Yuille A. Genetic CNN. In: Proceedings of the IEEE international conference on computer vision; 2017. p. 1379–88.

[127] Cai H, Zhu L, Han S. Proxylessnas: Direct Neural Architecture Search on Target Task and Hardware. arXiv. 2018.

[128] RUBICON. https://github.com/Xilinx/neuralArchitectureReshaping;.

[129] Hochreiter S. The Vanishing Gradient Problem During Learning Recurrent Neural Nets and Problem Solutions. International Journal of Uncertainty, Fuzziness and Knowledge-Based Systems. 1998;6(02):107–16.

[130] Graves A, Fernández S, Gomez F, Schmidhuber J. Connectionist Temporal Classification: Labelling Unsegmented Sequence Data with Recurrent Neural Networks. In: Proceedings of the 23rd international conference on Machine learning; 2006. p. 369–76.

[131] Bushnell, Brian. BBMap;. http://sourceforge.net/projects/bbmap/.

